# Reconstruction of TrkB complex assemblies and localizing antidepressant targets using Artificial Intelligence

**DOI:** 10.1101/2023.02.21.529454

**Authors:** Xufu Xiang, Chungen Qian, Hanbo Yao, Pengjie Li, Bangning Cheng, Daoshun Wei, Wenjun An, Yuming Lu, Ming Chu, Lanlan Wei, Bi-Feng Liu, Junfa Xu, Xin Liu, Fuzhen Xia

## Abstract

Since Major Depressive Disorder (MDD) represents a neurological pathology caused by inter-synaptic messaging errors, membrane receptors, the source of signal cascades, constitute appealing drugs targets. G protein-coupled receptors (GPCRs) and ion channel receptors chelated antidepressants (ADs) high-resolution architectures were reported to realize receptors physical mechanism and design prototype compounds with minimal side effects. Tyrosine kinase receptor 2 (TrkB), a receptor that directly modulates synaptic plasticity, has a finite three-dimensional chart due to its high molecular mass and intrinsically disordered regions (IDRs). Leveraging breakthroughs in deep learning, the meticulous architecture of TrkB was projected employing Alphfold 2 (AF2). Furthermore, the Alphafold Multimer algorithm (AF-M) models the coupling of intra- and extra-membrane topologies to chaperones: mBDNF, SHP2, Etc. Conjugating firmly dimeric transmembrane helix with novel compounds like 2R,6R-hydroxynorketamine (2R,6R-HNK) expands scopes of drug screening to encompass all coding sequences throughout genomes. The operational implementation of TrkB kinase-SHP2, PLCγ1, and SHC1 ensembles has paved the path for machine learning in which it can forecast structural transitions in the self-assembly and self-dissociation of molecules during trillions of cellular mechanisms. In silicon, the cornerstone of the alteration will be artificial intelligence (AI), empowering signal networks to operate at the atomic level and picosecond timescales.

## 1. Introduction

MDD, a systemic psychiatric disorder, may have a wide range of implicit etiologies: synaptic misconnection, metabolic abnormalities and immune inflammation. The lifetime prevalence of MDD surpasses 20 percent in the worldwide population, and the unavailability of specific medications renders one-third of patients unresponsive to treatment [1]. As the initial spot of the signal cascade, transmembrane receptors are indispensable in gaining and transferring signals across a thousand trillion interconnected synapses and as the objective of fifty percent of prescription medicine [2,3]. Several ADs have a direct link to the 5-hydroxytryptamine receprots (5-HTRs) of GPCRs and the ion channel-type glutamate receptors NMDAR and AMPAR for the onset of action [4]. Subtypes of 5-HTRs binding maps with ADs have been constructed then using virtual pharmacological screenings of millions of compounds to catch novel non-hallucinogenic antidepressants [5-9]. The results of conjugating NMDAR with S-ketamine reveal that S-ketamine takes effect more rapidly than conventional ADs by inhibiting Ca^2+^ influx to intracellular [10]. Previously, the predominant hypothesis claimed that ADs indirectly positively regulate TrkB via NMDAR to trigger the synaptic plasticity mechanism [11]. However, in a latest report, robust R-ketamine release therapeutic advantages by directly binding to TrkB [12]. This casts doubt on the broadly held 5-HT and NMDAR hypotheses and propels TrkB to spring up as a momentous priority receptor in building neat antidepressants [13].

Determining protein structure is essential for pharmaceutical research, while there is a deficit of architectural insights into TrkB despite the profusion of foundational research results [14-16]. TrkB modulates postsynaptic protein expression and synaptogenesis through the MAPK, PIK3/mTOR, and PLC pathways, intimately associated with a multitude of psychiatric conditions, including depression, Parkinson’s disease, and schizophrenia [17-20]. Like many other RTKs, due to their variable transmembrane topology and molecular mass outweighing the upper limit of conventional resolution approaches, only fifty percent of TrkB sequences have solved. Several RTKs, including EGFR, INSR, and ALK, have accurate measurements of the extracellular segment thanks to recent methodological enhancements in Cryo-electron microscopy (Cryo-EM) [21-23]. However, the membrane-spanning and cytoplasmic sections of all ligand-binding multimers were presented in low resolution due to IDRs [24-26]. The dearth of architectural information has sparked debate on a variety of hot-button issues, along with the following:

1. Why the extracellular TrkB segment reacts to distinct NGF-family members having diverse biological responses [27]?
2. Does TrkB’s transmembrane helix bridging angle alter the on/off state and strength of its subsequent enzymatic activity, and is it a feasible curative landing point [28,29]?
3. How does TrkB’s intracellular kinase element engage over 140 chaperones to assemble signal gatherings and automatically disentangle post-phosphorylation [30]?

Only a few layouts of RTKs bound to key partners were fixed since resolution entails a rigorous selection of docking sequences, kinase phosphorylation phase manipulation, and crystallographic strategies [31,32]. AF2, a deep learning-based artificial intelligence algorithm, predicted and released the structures of all protein sequences, making them easily accessible before resolution [33]. Previously, the G protein-coupled receptor (GPCR) and ATP-binding cassette receptors were correctly identified as the experimental structures adopting AF2 [34,35]. RTKs predicting topologies are rarely utilized because of their trisecting topologies and characterization of dimer activation [36]. As a result of AF2 advancements, biologists can construct biochemically active aggregates utilizing the protein complex prediction algorithm AF-M [37]. Even if TrkB is unresolved, AF-M can build its signal assemblies, and clear out if the TrkB dimeric transmembrane helix’s architecture does bind to ADs [33,38].

In the first step, utilizing AF2, the full-length monomer of TrkB was built with atomic-level precision. Due to IDR, the full-length dimeric structure of TrkB was not topology-compliant. To tackle this problem, the TrkB sequence was divided into extracellular, transmembrane, and intracellular segments. TrkB extracellular portion relating to mBDNF has predicted, and it is consistent with the resolved structure possessing the similar intrinsic and specific ligand-binding regions [15]. Creating the model of TrkC linking NTF3 indicates that AF-M rapidly pair between homologous receptors and ligands. The structural information help design microproteins with regulatory activity for a specific pair [39]. In physiological membrane environment, the transmembrane helix dimers maintained stable, and binding pocket for the novel ADs: IHCH-7086 and (2R,6R)-HNK was localized in the helix crossing point [9,40]. This discovery expands the list of pharmacological targets from GPCRs and ion channel-type receptors to include RTKs [41,42]. AI detected the essential posture of the assemblies during phosphorylation using related signal substrates like PLCγ1, SHP2, and SHC1 [11,30]. In conclusion, the TrkB’s snapshots during the signal cascade have been effectively reproduced using AF-M. Networks of physiological or pathological signal pathways comprising spatiotemporal information will be built in computers with artificial intelligence. In the next generation, AI will provide crucial points of the unresolved protein architectures associated with pathologies, broadening the scope of structure-based drug discovery to all coding sequences.

## 2. Results

### 2.1. AF2 predict the full-length and dimeric structures of TrkB

#### 2.1.1. Predicting the full-length structure of TrkB monomers by three methods using AF2

Previously, partial sequences of TrkB have been resolved by X-ray or nuclear magnetic resonance (NMR), including the ligand-binding segment (283-385) [43], the juxtamembrane IDR (497-519) [44], and the kinase segment (527-822) [16]. However, the structure of the remaining extracellular segment and the transmembrane helix structure is yet unknown. The full-length structure of TrkB predicted by AF2 was obtained in three ways:

4. Directly from the Alphafold Database (DB), which released structures in bulk in June last year based on Monomer V2.0 (Fig S1A);
5. Using Monomer V2.2 (Monomer), based on the adoption of the latest database, which entered the full-length sequence of TrkB from Ab initio prediction (Fig S1B);
6. Ab initio prediction using the Monomer Casp14 (Casp14) algorithm (Fig 1B), which consumes eight times the computational resources of Monomer but has higher accuracy.

The fixed structures predicted by the three modalities were similar, TrkB consists of five major structures, Domain 1: two CR clusters sandwiching three LRRs, Domain 2: Ig C1, Domain 3: Ig C2, Domain 4: transmembrane α-helix, and Domain 5: kinase domain (Fig 1A). However, the main difference between the predicted results could be attributed to the disordered sequences on both sides of the transmembrane helix and at the N and C-terminals. The PLDDT values of these sequences are shown to be low and as orange noodles by AF2. The disordered sequences are freely distributed, resulting in the predicted extracellular, transmembrane, and intracellular relative positions of TrkB not conforming to their spatial distribution and separated from each other at the plasma membrane.

**Fig 1.**
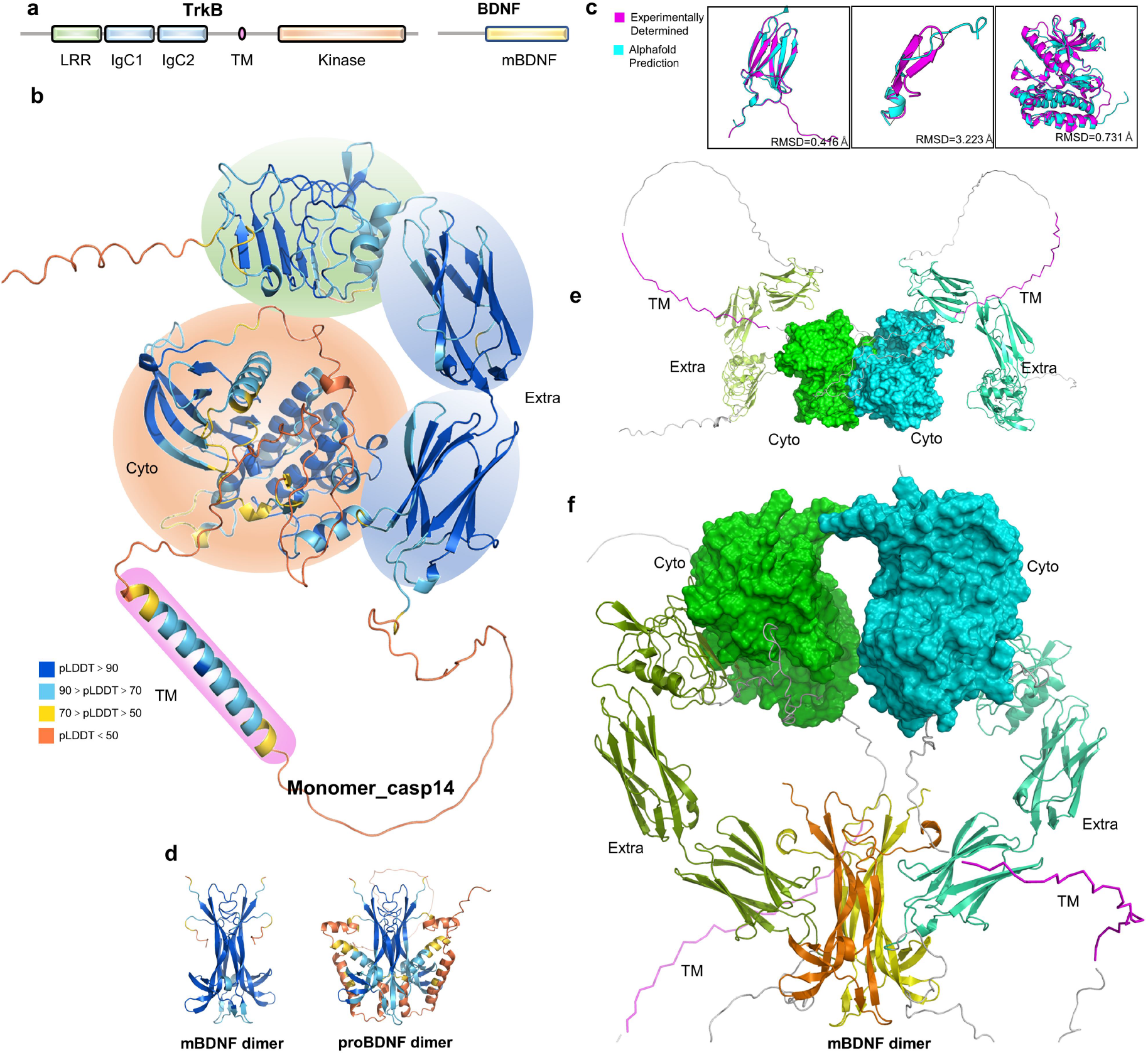
TrkB monomer, dimer structure predicted by AF2. (**A**) Schematic diagram of the structural domains of TrkB and BDNF. (**B**) The full-length structure of TrkB predicted by casp14, with different interval confidence levels displayed in the structure in the corresponding colors. Fixed structures are circled and colored in line with (**A**). (**C**) Structural alignment between the AF predicted structure and the experimentally resolved structure, in order: IgC2 [PDB:1WWB], juxtamembrane IDR [PDB:2MFQ], kinase domain [PDB:4ASZ], the RMSD between the predicted structure and the experimental structure is marked in the lower right corner, respectively. **(D)**mBDNF and proBDNF homodimeric structures predicted by AF-M. (**E**) The ligand-free dimeric structure of TrkB predicted by AF-M does not possess a reasonable membrane topology. (**F**) AF-M predicted structure of the activation dimer of TrkB-binding mBDNF.

The comparison of the PLDDT values of the predicted structures obtained by the three methods revealed two main differences (Fig S1D). Monomer’s transmembrane helix has a higher confidence level than DB, and the ends of the helix are extended (Fig S1F). The high confidence sequences coincide with the intersection of the transmembrane helix when it dimerizes, a site that has a fixed structure when dimerized to the ends and is captured by Monomer [45,46]. This discrepancy is due to DB augmentation or iterations of the algorithm, suggesting that although the DB already provides immediately available structures, ab initio predictions based on the most recent DB are necessary before the study. Another difference exists in the activation loop of the kinase, wherein the Casp14 algorithm has a 10% lower PLDDT value than the other two algorithms, and the activation loop shows a tightened and expanded conformation at different stages of kinase phosphorylation [47], overlapping the three predicted structures; also, a large difference was noted in the relative position of the activation loop (Fig S1C). This finding indicated that the Casp14 algorithm can identify such variable conformations during predictions compared to the Monomer algorithm, which confers a lower confidence level on the Casp14 algorithm.

#### 2.1.2. The fixed structure predicted by AF has atomic-level accuracy, while the disordered region is less accurate

The resolved structure files of TrkB were obtained from PDB, and the structures were compared to the predicted structures obtained by the three methods (Fig 1C and S1E). Strikingly, the predictions obtained for both Domain 3 and Domain 5 have a high prediction accuracy, with the Casp14 algorithm being better than DB in predicting Domain 3 and Domain 5, with an error reduction of about 0.2 Å, decreased to 0.416 Å and 0.731 Å, respectively. On the other hand, the RMSD of the predicted structure to the true structure reached about 3 Å when comparing the disordered region upstream of the kinase segment. Moreover, the sequence in the predicted structure behaves as an incomplete helix, while two tandem β strands are present in the resolved structure. The experimental structure was resolved by synthesizing a short peptide of this partial sequence and binding to the signal protein FR2Sα, which is a property of disordered sequences that can only be resolved as a fixed structure after binding to a specific signal protein [24,44]. Thus, AF2 has atomic-level accuracy in the prediction of fixed structure but does not provide an effective reference for the disordered region.

#### 2.1.3. AF-M predicts the full-length dimer structure of TrkB without ligand/binding ligand and predicts the spatial distribution of the structure irrationally

AF-Multimer, used for protein complex prediction, has been released recently and has demonstrated excellent performance in constructing dimers and multimers [24,48]. It has been used for dimer construction of TrkB to understand the usability of AF2 for RTKs, which have dynamic conformations and a wide range of binding partners and complex binding forms [36]. The structure of TrkB has not yet been resolved at high resolution under Cryo-EM, and AF2 does not allow direct access to experimental dimer structures. Nevertheless, the extracellular dimer and transmembrane dimer of the TrkB homologous protein TrkA have been resolved to use it as a reference for the predicted structure of TrkB to evaluate the availability of AF-M predictions [49,50].

Previous studies have suggested that TrkB has a ligand-free dimeric structure, and the full-length sequences of TrkB were set as sequence 1 and sequence 2 and input into AF-M for prediction [51,52]. Among the first five results obtained, the dimer contact interfaces of those ranked 1– 4 exist between the kinase segments, and no contact exists between the two monomers of that ranked 5, while the extracellular and transmembrane segments does not possess any reasonable relative space, and the main difference between the structures is the spatial distribution of the extracellular segments (Fig S2A). For example, in ranked 1, the contact occurs mainly on the C-terminal of the kinase segment, while the extracellular segment bends outward near the kinase of the same chain, and the transmembrane helix loses its helical conformation and is recognized as IDR and wrapped around the outer side (Fig 1E).

To rescue this failed spatial conformation, an attempt was made to add its high-affinity ligand BDNF for the prediction of TrkB dimer (Fig 1D and S2D) [27]. First, it was essential to demonstrate whether AF-M could successfully construct BDNF dimers, which exist in two forms during secretion, including precursor BDNF (proBDNF) at full length, and mature BDNF (mBDNF), enzymatically cleaved at the N-terminal. The binding of proBDNF to p75 mediates long-lasting inhibition and synaptic disappearance, whereas binding of mBDNF to TrkB enhances synaptic plasticity and synapse formation [11,53,54].

The homodimers of the two forms of BDNF were constructed independently (Fig 1D and S2D). mBDNF dimer is highly similar in structure to the already resolved mBDNF/NTF-4 heterodimer [55]. The main body of mBDNF consists of three pairs of β-strands and four linked β-hairpin loops, and the dimer is centrosymmetric with the long axes of two pairs of long β-strands contact (Fig 1D and S2D). In contrast, the structure of the proBDNF dimer has not yet been resolved, and the predicted structure has four incomplete helices at the N-terminal of each monomer. The presence of these helices caused the lower end of the β-strands to be pulled to both sides, which disrupted the intrinsic binding interface of the NGF-β family at the IgC2 of the Trk family (Fig S2C). This alteration revealed the reason for the low affinity of proBDNF to TrkB and high affinity to p75, providing a structural explanation for the opposite synaptic effect of proBDNF and mBDNF [27,56]. Consequently, AF predicted mBDNF homodimers, demonstrating that it is highly usable in the prediction of dimeric ligands and can rapidly predict all the putative intrafamily paired dimer conformations when homologous family dimeric structures are available.

Based on the successful construction of mBDNF dimer, we further predicted the conformation of TrkB full-length dimeric activation state, TrkB-mBDNF-mBDNF-TrkB (Fig S2B). The major dimerization interface of all predicted results was the extracellular segment, and the kinase dimerization structure in the ligand-free dimerization structure was separated, indicating centrosymmetric but no contact. In ranked 1 as an example: mBDNF bridges the bipartite structure of the extracellular segment, while the kinase segment is incorrectly placed at the top of the N-terminal of the extracellular segment and does not contact each other (Fig 1F). The transmembrane helix is recognized as a disordered structure, indicating that the addition of mBDNF does not rescue the irrational spatial distribution of TrkB dimeric conformation predicted by AF-M.

In conclusion, we found that AF-M does not provide a reasonable full-length dimeric structure of TrkB with or without binding ligands. This phenomenon is attributed to the fact that the resolution results of RTKs Cryo-EM could not provide a sufficient density for the transmembrane and intracellular segments due to the lack of rigid connections between the extracellular segment and transmembrane helix, and only the dimeric extracellular segment was resolved at high resolution and uploaded to the database [20]. Despite the inability to give a full-length dimeric structure, AF-M provides a dimeric conformation of the kinase segment and the extracellular segment of the bound ligand, derived from the mimicry of the local structures of RTKs obtained by NMR and X-ray over the past 30 years. This indicates that AF-M has the ability to predict the fixed structure of RTKs undergoing dimerization. Hence, we split the sequence of TrkB into extracellular/transmembrane helix/intracellular segments for dimerization prediction based on previous studies of EGFR and predicted their complex conformation upon binding ligands, signal components, or drugs [28].

### 2.2 AF-M accurately predicts the ligand-binding structure of the extracellular segment of the TrkB dimer

#### 2.2.1 Prediction of potential TrkB extracellular ligand-free binding dimer structure with inter-monomeric binding via β-turn

The predicted TrkB extracellular ligand-free dimer was obtained by AF-M. The docking of ranked 1–ranked 3 occurs at Domain 3, while the difference between the structures is that the two monomers Domain 3 are increasingly spaced apart with decreasing confidence until no contact is made; the docking of ranked 4-5 occurs at Domain 1 and IDR (Fig S5A). These findings differed from the previously experimentally obtained dimeric structure of Trk family Domain 3, wherein the crystal structure is based on the two monomers with overlapping N-terminus and C-terminus, which results in the loss of βA at the N-terminus of Domain 3 [57]. Such a structure is considered erroneous because Domain 2 exists on Domain 3 N-terminal, and the N-terminal sequence serves as a bridge between Domain 2 with 3. Despite the incorrectly constructed Domain 3 dimer in the database, AF-M can determine and correct its implausibility. In ranked 1, βA was successfully identified and linked to Domain 2, and the monomers contacted each other through the loop between βA and βB (ABL), i.e., the interaction force of S297-W301 (Fig 2A and 2B). A total of five hydrogen bonds and two hydrophobic bonds were formed between the residues, and a π-π stacking was formed between H299 and W301. These residues formed a negatively charged finger-like protrusion and a concavity, and the two monomers formed a chimeric structure with each other, which was consistent with the subsequent prediction of a specific ligand-binding region (Fig 2C). Furthermore, M379 and G380 near the membrane were also identified as bind residues in the disordered region.

**Fig 2.**
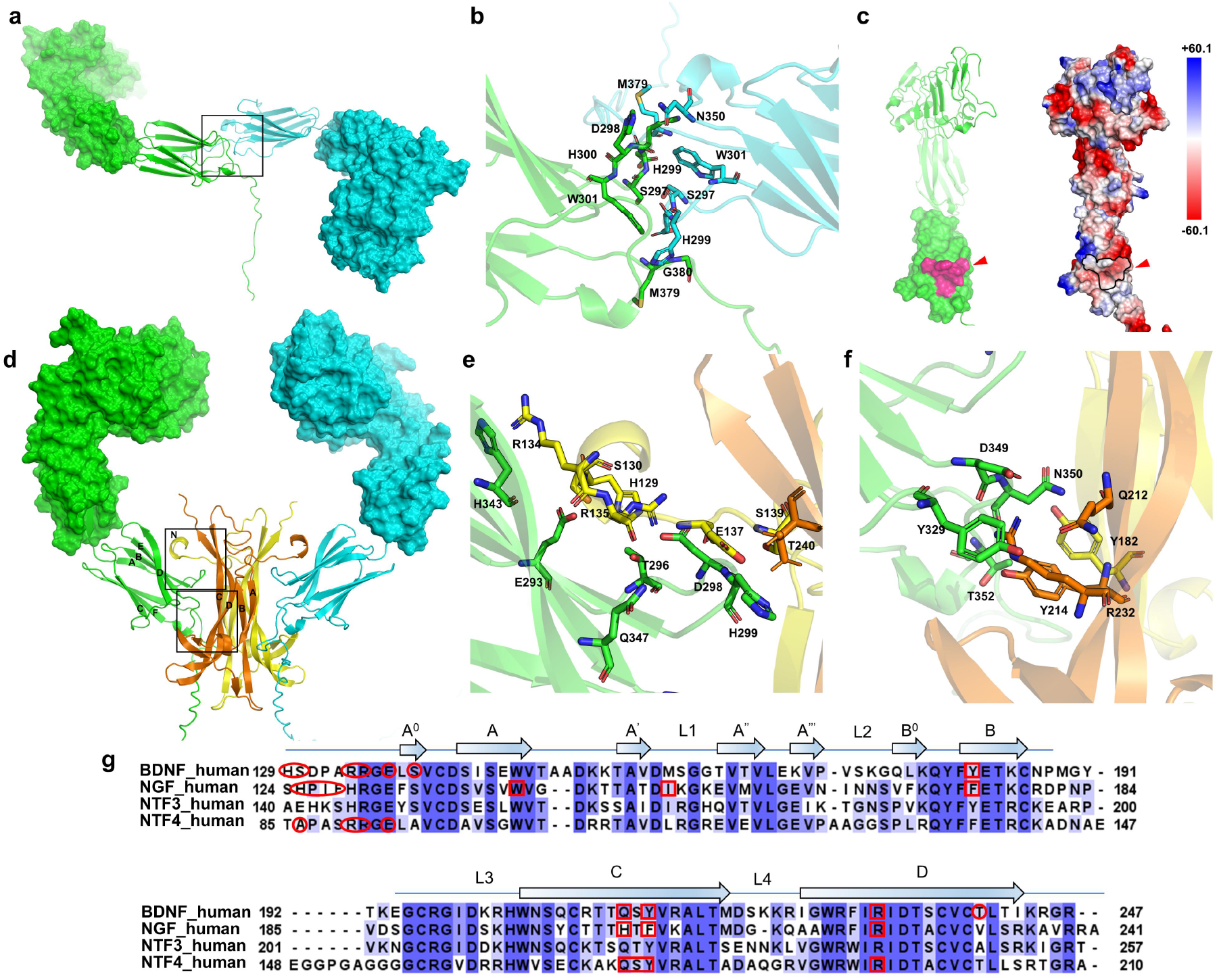
Structure prediction of pre-activated and activated dimer of TrkB extracellular segment. (**A**) AF-M prediction of the potential TrkB extracellular segment pre-activated dimer structureb. (**B**) Detailed graphical representation of the contact residues of the pre-activated TrkB pre-dimer. (**C**) TrkB pre-dimer contact residues form finger-like protrusion and concavity, marked in purple (left). Circled in black stroke on the surface potential energy map (right). (**D**) AF-M prediction of the extracellular activation state dimer of TrkB that binds mBDNF. The two interact regions are framed in black. (**E**) Residue details of the TrkB-binding mBDNF-specific binding region are shown. (**F**) Detail map of residues in the intrinsic binding region of TrkB-binding mBDNF is shown. (**G**) Sequence alignment of NGF-β family, where residues in the specific binding region are circled and residues in the intrinsic binding region are boxed. Information of BDNF is from the predicted structure of TrkB-mBDNF. Information of NGF NTF4 is from the resolved structure of TrkA-NGF and TrkB-NTF4.

AF-M offers the possibility of an unresolved TrkB pre-dimer, i.e., ABLs that contact Domain 3 specifically with ligands forming chimeras with each other, and the ligand-binding surface is buried until the ligand is inserted and opened to stimulate the downstream signals.

#### 2.2.2 Predicting the ligand-bound dimeric structure of TrkB extracellularly with specific binding region and intrinsic binding region between Domain 3 and mBDNF

Next, AF-M was used to predict the conformation of the extracellular segment of TrkB bound to mBDNF (Fig S4A). The five results showed a high degree of agreement, with differences arising from the free distribution of the disordered region. The complex structure of mBDNF retains its centrosymmetric homodimeric conformation and binds to the Domain3 of TrkB. Conversely, the extracellular segment of TrkB has a crab-pincer shape with two monomers each monomer is attached to a single chain, alternating between the front and back of mBDNF. This finding is similar to the results of the TrkA extracellular segment [58]. The most different secondary structure is Domain 1, wherein β1–β3 starting from the N terminus is shorter in the crystalline structure than in the predicted structure, and most of the articulated sequences have a loop-like morphology (Fig S5B). These results predicted a TrkB Domain 1 superhelical topology compared to the resolved TrkA.

Taken together, the difference between the real structure of TrkA and the predicted structure of TrkB lies in the relative positions of Domain 1–Domain 3. Compared to TrkB, located on the same face as Domain 1–3, TrkA produces a large-angle bend between Domain 1–3, which is nearly Z-shaped from the side (Fig 2D and 2G). This is because the de-glycosylation strategy is avoided during the resolution of TrkA. Asparagine-linked glycosylation plays a key role in the regulatory activity, and the six glycosylation sites are highly conserved in the Trk family, located at the N-terminal and C-terminal ends of Domain 2, bridging Domain 1 and Domain 3 [58]. On the other hand, AF-M ignores this post-translational modification (PTM), allowing the arms stretched tightly by glycosylation to be predicted in a relaxed conformation. The structure of the ligand-binding Domain 3 is highly consistent with the resolved TrkA-NGF and TrkB-NTF4 structures [15,59]. The sequence comparison of NGF-β family revealed two main binding regions between Trk and NGF-β families: specific binding region and intrinsic binding region (Fig 2E). The specific binding region exists in the highly variable N-terminal sequence of each NGF-β family member, while the intrinsic binding region is found in βC and βD; both are highly conserved across all NGF-β family members as the backbone structure.

Specifically, the specific binding region is located in the ABL of TrkB Domain 5, which is warped and embedded in the groove formed by the N-terminal end of mBDNF (Fig 2F). TrkB-contacting residues are located in ABLs: E293, T296, D298, and H299; βE: H343 and Q347. A total of nine hydrogen bonds were formed between the chains, and the most tightly bound residue was H299, on which three hydrogen bonds were generated. In the case of mBDNF, the residue in contact with TrkB is located at the N-terminal H129–S139, where the structure shifts from a disordered state when unbound to an incomplete helix. This indicated that AF predicts the transition from the disordered to a fixed secondary structure upon binding of the ligand to a specific receptor. Moreover, the intrinsic binding region is present on mBDNF at the intersection of βC and βD, including Q212, Y214, R232, and R182 of βB. These residues are highly conserved in the same family, while the contact residues on TrkB Domain 5 are D349, N350, and T352 of EFL; the tightest connection occurs between D349 and R232, forming two hydrogen bonds.

To further verify the usability of AF-M in predicting the ligand-receptor binding structure, most of the sequences of the extracellular segment of TrkB was deleted, and only Domain 3 was retained and the tetramer with mBDNF was reconstructed (Fig S4C). The obtained results were consistent with those obtained with the whole extracellular segment. Subsequently, the homologous sequence of NTF3 and TrkC was selected for construction (Fig S4D). Although only the monomeric Domain 3 was previously resolved by TrkC, the dimeric ligand-binding structure was obtained successfully by AF-M [43]. This finding indicated that AF2 can predict the structure of complexes with ligands encompassing minimal ligand-binding structural domain sequences, as well as various pairing structures of homologous family receptors with ligands.

In conclusion, in the structure prediction of extracellular segment-binding ligands, AF-M is unable to predict the effect of PTMs on the structure, resulting in a relative angle between domains deviating from the true structure. However, the predicted structures are highly accurate for the ligand-bound receptor structural domains, constructed with reference to the experimental structures between ligand-same family receptor members of the same family in the database. Previous studies have shown that several RTKs can form heterodimeric pairings with homologous receptors [55,60,61]. Also, ligand family members can heterodimerize, with dozens of possible pairings between the Trk family and the NGF-β family alone. AF-M facilitates the assembly of all the ligand-receptor binding structures using only the sequences, thereby circumventing the limitation of the resolved structures. Thus, we can retrieve highly matched design structure microproteins and small-molecule drugs using the three-dimensional structural information of proteins, thereby minimizing the side effects of the traditional antibody binding to the same family of receptors and regulating the activation and inactivation of receptor kinases at the atomic level [62,63].

### 2.3 AF-M predicts the antidepressant binding pocket at the crossover of TrkB transmembrane segment dimer

#### 2.3.1 The predicted TrkB transmembrane segment dimer structure is significantly different from the resolved structure of TrkA

The transmembrane helix dimer of TrkB was successfully constructed by AF-M (Fig S6A). The transmembrane helix dimerization structure of TrkA has been resolved previously [50].The predicted transmembrane helix crossover pattern of TrkB is significantly different from that of TrkA, with TrkA monomers further apart from each other and the overall structure similar to X-type. On the other hand, the structure of TrkB tends to be parallel, and the helix of TrkB is longer than that of TrkA, with the crossover site ASVVG located at the center of the helix, while the crossover site of TrkA is near the N-terminal (Fig S6B and S6C) [45]. Sequence comparison revealed that the crossover site SXAVG of TrkA and TrkC was shifted up by 7 residues compared to TrkB, indicating that the crossover site of TrkB is highly specific in the Trk family (Fig 3D). The sequence comparison of TrkB between different species revealed that the transmembrane helix is conserved in the family. While the transmembrane helix is a common drug binding site for GPCRs and ion channel-type receptors, the complex formation structures of various antidepressants with the 5HT family and the glutamate receptor family have been resolved [5,10,64,65]. Previous studies on EGFR have shown that the transmembrane helix transmits signals from the extracellular segment to the intracellular segment by changing the rotation angle and docking site [28,29]. This alteration implies that the transmembrane helix of TrkB has the potential to modulate the activation strength of the intracellular kinases and becomes a target structure for antidepressants.

**Fig 3.**
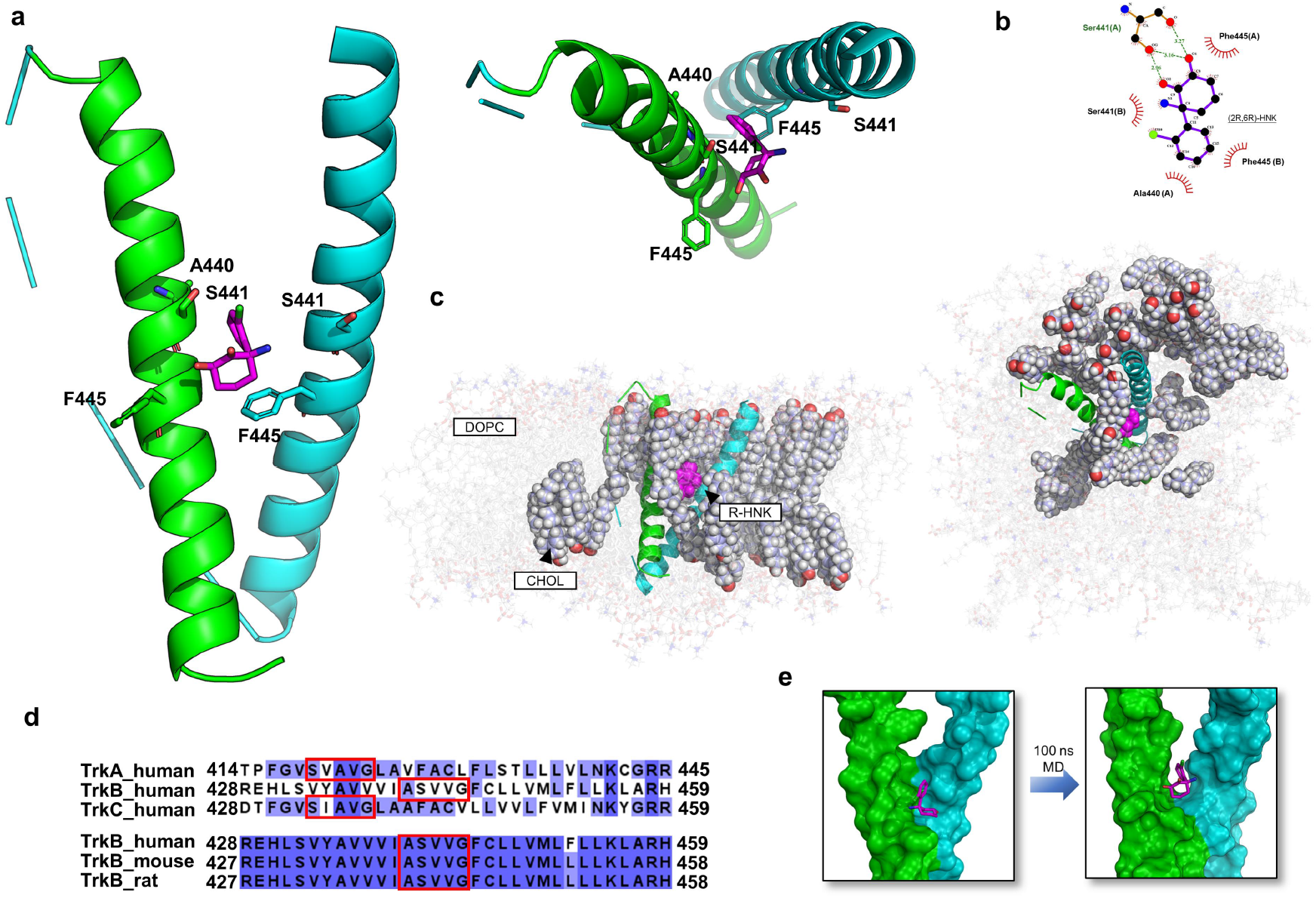
AF-M predicted TrkB transmembrane helix dimer after molecular dynamics simulation to localize the drug binding pocket (**A**) TrkB transmembrane helical dimer with (2R,6R)-HNK coupled structure after 100 ns MDS. Contact residues are shown in stick form. The top view is shown on the right. (**B**) Pose fingerprint of the (2R,6R)-HNK contact with the surrounding TrkB transmembrane helix residues. The most stable contact occurs on Ser441, with a total of three hydrogen bonding contacts. (**C**) The final conformation of the TrkB transmembrane helix dimer with the (2R,6R)-HNK-coupled structure in the lipid box, with cholesterol being displayed in a spherical shape, after 100 ns MDS simulation, the top view(right). (**D**) Sequence comparison of the human Trk family (top), and sequence comparison of TrkB across species (bottom), reveals that the crossover sequence is highly specific in the same family and highly conserved between species, and the helical crossover residue sequence is boxed in red. (**E**) After 100 ns MDS simulation, (2R,6R)-HNK is transferred from the protein surface to the crossover gap and a drug binding pocket is generated.

#### 2.3.2 Molecular dynamics simulations reveal a potential antidepressant binding pocket at TrkB docking

Previous studies used protein docking to obtain five possible transmembrane helix conformations based on the predicted monomeric structures and simulated them in the membrane environment by molecular dynamics [12]. Finally, only one docking could stabilize the dimerization in the membrane environment, and the conventional drug fluoxetine could bind to this site. The unstable dimerized helix could be ascribed to the fact that the prediction of protein docking for transmembrane helix does not consider the membrane environment and can only be screened by MDS on a large number of potential structures, which assigns a high threshold to the study of RTKs’ dimerization helix with drug docking.

The transmembrane helix structures predicted by AF-M were derived from the imitation of similar structures in the database. The AF-predicted transmembrane helix structures have been demonstrated to be highly accurate based on the comparison between the GPCR and ABC proteins with the resolved structures [34,66]. The five TrkB transmembrane helix dimerization structures predicted by AF-M differed only in relative angles, with consistent crossover sites, indicating that TrkB dimers are formed in cholesterol-rich lipid rafts [67]. To demonstrate the stability of the predicted structures in the membrane environment, the predicted ranked 1 dimer helix was placed in a 20% CHOL+80% DOPC environmental box, and molecular dynamic simulations were performed for 100 ns, and the dimer structures were found to be cross-stable until the end of simulation.

Two antidepressants, (2R,6R)-HNK and IHCH-7086, were selected for molecular docking with ranked 1 [68]. (2R,6R)-HNK exerts a rapid antidepressant effect via a non-glutamate receptor-dependent BDNF/TrkB pathway, while IHCH-7086 is a synthetic without psychedelic group derivative of Psilocybin derivatives that acts on the 5-HT receptor [9]. Both drugs contact the TrkB transmembrane helix near the crossover at V437-F445, indicating a potential drug binding site (Fig S7G and S7H).

In order to obtain the interaction of the drug with the transmembrane helix in the physiological membrane environment, the (2R,6R)-HNK docked dimeric transmembrane helix structure was subjected to MDS in 20% CHOL+80% DOPC environment (Fig 3C). We found that the transmembrane helix showed a tendency to converge and reach a steady state in the second half of the simulation. The RMSD values fluctuate from 0–50 ns, which is caused by the initial instability of the docked acquired conformation inside the box, which declines rapidly at 50 ns and enters the steady state at 60 ns. The fluctuation of Rg corresponds to the ripple correspondence in RMSD, wherein the fluctuation rises in the first period, has a small peak at 10 ns, starts to reach a second peak at 50 ns and finally enters the stable zone at 75–100 ns, indicating that the system is shifting from the unstable to the stable state at this moment. The SASA of the protein gradually decreases in 0–100ns, indicating a favorable binding and gradual protein tightening. HBNUM showed 0–2 connections of hydrogen bonding during the simulation (Fig S7A-S7D).

Analysis of the variation in the binding free energy with MDS between protein-ligand complexes showed a total free energy of -17.14 kcal/mol, indicating a high probability of interaction between TrkB and (2R,6R)-HNK (Fig S7E and S7F). After breaking down each subterm, VDWAALS <0 indicates that hydrophobic interactions contribute to binding, and EEL <0 indicates that out hydrogen bonding also contributes to binding. EGB >0 indicates that polar solubilization is not conducive to binding, while ESURF <0 indicates that nonpolar interactions are conducive to binding.

Postural fingerprinting of the stable conformation revealed that the interaction of (2R,6R)-HNK with the dimeric transmembrane helix originates from hydrogen bonding and the surrounding hydrophobic amino acids, whereby (2R,6R)-HNK produced hydrogen bonding with S441 and hydrophobic forces with V437, V438, A440, V442, and F445 (Fig 3A and S7g). Further analysis of drug-residue interactions revealed three hydrogen bonds created between (2R,6R)-HNK and S441 at a distance of about 3 Å (Fig 3B). The comparing of (2R,6R)-HNK before and after the simulation showed that the molecule moved from the protein surface to the center of the crossover, where it formed a TrkB-specific hydrophobic pocket that was exactly on the two dimer crossover sequences (Fig 3E). This phenomenon suggested that (2R,6R)-HNK effectuates the extracellular segment signal on the kinase segment by anchoring the transmembrane helical dimer conformation, thereby exerting a synaptic plasticity and enhancing the antidepressant effect.

The structural information of the transmembrane segments of RTKs obtained by conventional methods is sparse because of the difficulties in sample preparation [50]. However, the crystallization-independent nature of Cryo-EM provides the overall dimeric conformation of RTKs and sets up different plasma membrane mimics to capture the dynamic conformation. Nevertheless, the transmembrane segments are in low-resolution spherical shape due to the lack of rigid articulation between the extracellular and transmembrane segments [20]. AF-M provides a stable dimeric transmembrane helix in the membrane environment and thus a reference structure for image acquisition and model reconstruction during Cryo-EM resolution, further improving the resolution [69-71]. Thus, the discovery and design of drugs targeting dimeric transmembrane helices has broadened the scope of drug research that previously targeted GPCR and ion channels.

### 2.4 AF-M predicts intracellular signal assemblies of TrkB with multiple binding modes between kinase and signal proteins

#### 2.4.1 AF predicts TrkB kinase homodimers with centrosymmetry between monomers centered on the activation loop

The kinase segments of the RTK family are structurally similar, with the main structure consisting of an N-lobe composed of α-helixes and a C-lobe composed of β-folds, as well as a hinge bridging the two globes, while a loop sequence, known as the activation loop, exists on the C-lobe and plays a critical role in the phosphorylation cascade [72]. For TrkB, there are five majors phosphorylated tyrosines (pTyr), including Y516 in the proximal membrane region recognized by AF as an incomplete helix, Y702, Y706, Y707 in the activation loop, and Y817 in the C-terminal IDR tail. The predicted TrkB kinase segment shows a consistent conformation with the resolved kinase-inactive structure, wherein the αC helix opens outward and the expanded binding pocket appears as C-helix out; the benzene ring of Phe in the DFG at β8 appears as DFG-out towards the hinge domain, indicating that the protein is inactivated and is in a closed state. Despite the presence of the DFG-in structure of CPD5N [PDB:4AT3] with the antagonist in the database, AF2 tends to select the inactive state of the kinase for output. Among all subsequent structures of the complex, the kinase presents an inactive conformation consistent with the monomer (Fig 4B).

**Fig 4.**
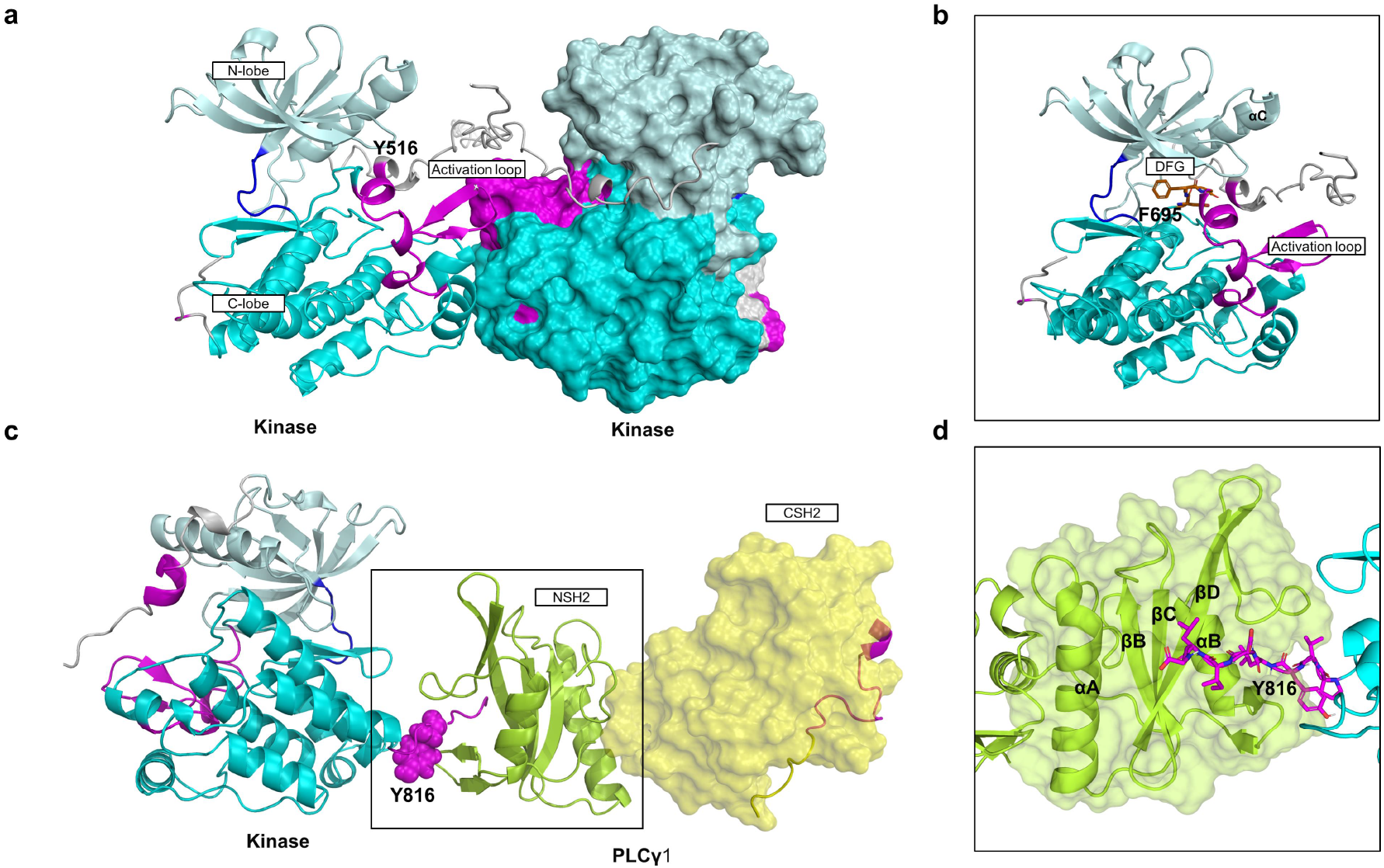
AF-M predicted TrkB intracellular kinase segment homodimer, heterodimer bound to PLCγ1 NSH2-CSH2 (**A**) AF-M predicted homodimer of TrkB kinase segment. The two monomers are centrosymmetric with the active ring as the central point, and the active ring is close but not in contact. The proximal membrane region crosses the interface where the active ring is located. The hinge region is dark blue, and the proximal membrane region containing the autophosphorylated tyrosine, the active loop, and the C-terminal tail are purple. Same below. (**B**) AF-M predicted sequences related to kinase activity in TrkB kinase segment homodimers are shown. αC is C-helix out, DFG is DFG-out, and the kinase is in the off state. (**C**) AF-M predicted heterodimer of TrkB kinase segment with PLCγ1 NSH2-CSH2 structural domain, Y816 site in contact with NSH2, shown as spherical. (**D**) Details of the contact of the TrkB kinase segment with the PLCγ1 NSH2 structural domain are shown the C-terminal tail is embedded in the surface groove of NSH2 and is localized near αB.

Furthermore, three models of self-inhibition have been identified in studies of RTKs, including cis-inhibition of the active site by the proximal membrane region and C-terminal tail, tightening of the activation loop, and blocking of the substrate binding site [72]. In the predicted structure of the TrkB monomer, the juxtamembrane region is identified as a disordered scattered structure, the C-terminal tail has only 10 residues and does not block the ATP-binding site, and the activation loop shows an outwardly expanded relaxed structure. Therefore, AF did not provide a monomeric TrkB self-suppressive structure.

The prediction of the dimeric structure of the intracellular segment of TrkB was attempted to understand the predictive power of AF-M for the dimeric conformation of the kinase segment. A previous electron microscopic analysis of EGFR and KIT found that the kinase existed in multiple dimeric conformations, showing asymmetric distribution in most states [25,26]. Conversely, the kinase dimers obtained by AF-M were symmetric in the center of the activation loop, which comes from the main principle of AF-M in predicting the homologous dimers, i.e., structural symmetry (Fig 4A and S8). This finding suggested that AF-M is unable to predict the dynamic conformation of the pTyr intercontact between monomers at the kinase dimerization stage, but identifies the central role of the activation loop in trans-phosphorylation. This phenomenon occurs at the proximal membrane end consisting of Y516. Among all the predicted structures, juxtamembrane sequence, which is originally free in space, crosses from the interface at the activation loop. This indicates a critical role of AF-M via juxtamembrane segment in coupled phosphorylation, wherein a contact was established between the proximal membrane segment and the activation loop during cascade phosphorylation.

#### 2.4.2 Predicted TrkB kinase segment complex structure with PLCγ1 and SH2 structural domain bound to the kinase C-terminus

Tyrosine kinases transmit signals downstream by recruiting and activating substrate proteins. pTyr binds to the SH2 and PTB structural domains of the signal proteins to generate cascade phosphorylation. The SH2 and PTB structural domains are present in a diverse set of proteins containing a range of catalytic type and interacting domains, which provide a degree of specificity by recognizing pTyr residues and the surrounding residues. Three extensively studied signal proteins were selected for the construction of their complex structures with TrkB kinase segments to examine the efficacy of AF-M in predicting the intracellular molecular assemblies of tyrosine kinases. Notably, the AF was previously shown to be effective in the prediction of intracellular giant complexes, with quiescent structures [71]. However, the tyrosine kinase signal is transmitted by a liquid-liquid phase separation mechanism, and the protein structure in the droplet exhibits a high degree of dynamics [38,73].

The PLCγ signal pathway is downstream of TrkB, which plays a major role in promoting calcium inward flow [74]. The activation of the pathway leads stimulates CaMKII and subsequent synaptic plasticity, where PLCγ1 serves as the first substrate with two tandem SH2 domains. The previous structural analysis provided two reference binding modes; one from the binding of NSH2-CSH2 to FGFR1, in which NSH2 forms a binding pocket on pTyr at the C-terminus of the kinase [31], and the other is from the complex formed by FGFR2 and CSH2 [75]. Based on this structure, another activation model was proposed as follows: CSH2 contacts both kinases simultaneously, and in addition to producing a pocket binding mode similar to that described above, the pTyr-containing tail at the C-terminus of CSH2 is inserted into the active groove of the other kinase. These two models provide insights into two types of kinase-substrate binding: 1. Substrate recruitment and phosphorylation are cis at the same tyrosine kinase monomer. 2. Substrate recruitment and phosphorylation are dependent on the dimerization of the kinase segment. Consequently, we selected NSH2-CSH2 of PLCγ1 and retained the pTyr-containing sequence downstream of CSH2 to construct the complexes of NSH2-CSH2 with TrkB kinase segment in a 1:1 ratio and CSH2 with kinase segment in a 1:2 ratio.

In a 1:1 complex, AF-M accurately predicts the attachment site for the cis reaction of monomeric tyrosinase with PLCγ1 (Fig S9A and S9B). The Y817-containing C-terminal tail of ranked 1–ranked 4 kinase is attached to NSH2, but the difference between the structures is due to the free C-terminal tail that points in a different direction. For example, in ranked 1, the C-terminal tail extends into the groove formed by NSH2 αB and βD. The tightest connection is between L821 and G822, wherein L821 is hydrogen bonded to βD H607 closest to the surface, while G822 is bonded to R562 on αA by two hydrogen bonds (Fig 4C and 4D). Although this interaction is on the same surface as the resolved structure, the penetration of the C-terminal tail of the predicted structure into NSH2 is in the opposite direction to the real structure, penetrating from the outside of αB and terminating at βD, while the contact of the resolved structure starts at αA (Fig S9C). The resolved structure of CSH2, which does not make contact with the kinase, is distinguished from NSH2 by AF-M and is located free in space despite the secondary structure similar to that of NSH2.

This finding indicated that AF-M predicted the signal chaperone contact domain and located the phosphorylation site in the case where the database contains the structure of the kinase complex with the signal companion. However, some differences were noted in the spatial distribution of specific residues from the resolved structure owing to sequence differences among the kinases and the additional possibilities offered by the database containing the pTyr peptide in a complex with the SH2 structural domain. Moreover, in the 1:2 complex, AF did not construct the theoretical model of transdimeric activation (Fig S10). CSH2 in all predicted structures was free from the dimerization of the kinase segment, indicating that despite the existence of the theoretical model, AF-M could not construct an approximate linkage due to the lack of reference structures in the database.

#### 2.4.3 Prediction of the complex structure of TrkB kinase segment with SHP2 and SHC1 and identification of the different binding modes of kinase binding to SHP2 disordered region and SHC1 PTB

To further understand the effect of AF on the formation of kinase-signal protein complexes, two key signal proteins were selected to construct the dimer formed with kinase, including SHP2 and SHC1 [76]. SHP2 is an allosteric enzyme that consists of two tandem SH2 structural domains, phosphatase structural domain and C-terminal disordered tail, while NSH2 contacts with PTP and self-inhibits the phosphatase activity in the absence of biochemical reactions [77]. In addition, SHC1 is a signal scaffold protein, and its PTB binds to the kinase structural domain, while SH2 further binds other signal proteins and large segments of the sequence show a disordered structure [30,78]. Different from PLCγ1, neither SHP2 nor SHC1 obtained a resolved structure of its structural domains in a complex with the kinase segment, and the inferred contact site was derived from the contact conformation of the pTyr-containing polypeptide with SH2 and PTB.

The predicted structure of the TrkB kinase segment with SHP2 does not provide a conformation in which the PTP is activated by conformational change; however, it still adopts a self-inhibitory structure, and NSH2-CSH2 does not interact with the kinase (Fig 5A and S11). A phosphorylated peptide study showed that the spaced pTyr site sequentially crosses the surface of SH2 and causes NSH2 to lose the inhibition of the phosphatase; then, the kinase contacts and reacts with the main body of the phosphatase as the conformation during the allosteric activation is not available due to the absence of PTM information from AF-M. The contact in the predicted structure occurs in the C-terminal α-helix of the PTP structural domain of SHP2 and downstream to IDR, as it is embedded in the active groove of the kinase forming a multivalent and stable contact, where Y547 of SHP2, the phosphorylated site creates four hydrophobic bonds and one hydrogen bond between L595, I692, and V673 of the kinase segment (Fig 5B and S11C). This interaction exhibits a transient local contact where SHP2 is recruited to the C-lobe as well as near the activation loop through the disordered region in the case of autoinhibition. Of the five possible conformations, four of the resulting contacts occur in the C-terminal disordered tail of SHP2. This interaction is consistent with recent studies on the liquid-liquid phase separation of SHP2, which undergoes LLPS in the autoinhibited state and is enhanced by the transformation of the phosphatase to an active conformation after mutation [77]. Before this phenomenon, AF2-based phase separation prediction programs have been developed by scoring disordered regions, while AF-M has the potential to complete the screening and localization of phase separation targets [79].

**Fig 5.**
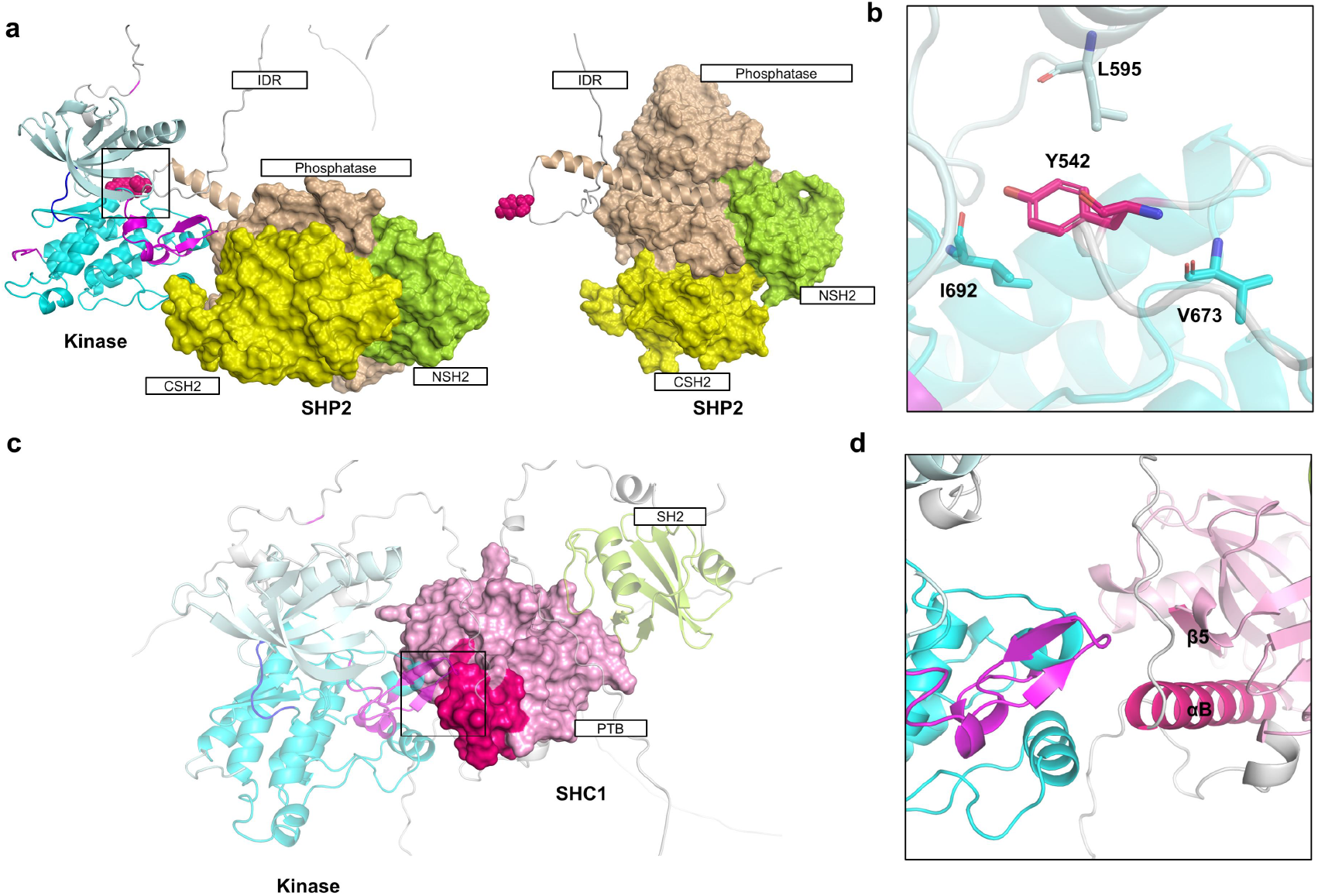
AF-M predicted heterodimer of TrkB intracellular kinase segment bound to SHP2 and SHC1 (A) AF-M predicted heterodimer of TrkB intracellular kinase segment SHP2, with the structure of the disordered region of SHP2 extending into the active groove between the kinase globes. (B) Detail of AF-M predicted heterodimer of TrkB intracellular kinase segment SHP2 showing that autophosphorylated Y542 on the disordered region of SHP2 makes contact with I692, V673 and L595 of αC of C-lobe. (C) AF-M predicted heterodimerization of the TrkB intracellular kinase segment SHC1 with contact occurring in the disordered region while PTB is closer to the C-lobe compared to SH2. No contact occurs between PTB and the kinase, but the binding pocket of PTB (red) is toward the C-lobe. (D)Detail of AF-M predicted heterodimerization of the TrkB intracellular kinase segment SHC1, with β5 and αB of PTB producing binding pockets toward the C-lobe and close to the activation loop.

In the predicted structure of the TrkB kinase segment with SHC1, the contact between the two occurs at the N-terminal IDR of SHC1 (Fig 5C and S12). Despite the inability to form an exact binding structure similar to that of PLCγ1, AF2 identified PTB as the structural domain of the SHC1-binding kinase. Consequently, PTB shows a spatial localization close to the C-lobe, SH2 is free on the outside, and the conserved binding pocket of PTB consisting of αB and β5 is opposite to the protrusion of the C-lobe (Fig 5D). This structure suggested that when confronted with unresolved fixed structure-bound kinase-signal protein complexes, AF2 can determine the structural domains that may be in contact, albeit without providing the exact docking residues.

In conclusion, we explored the potential conformation of AF-M in predicting the intracellular kinase segment dimerization and kinase segment binding to various key signal proteins. In the case of resolved structures with similar patterns, AF-M could accurately localize to the binding residues, such as NSH2 of PLCγ1. However, the monomer in the unresolved complex exhibits a self-inhibited structure with the blocked binding interface, and the predicted contacts occur at the IDR because the PTM information is not considered. In order to construct a complete, signal pathway network containing three-dimensional information is necessary to deduce additional conformations of kinases and companions at different reaction stages and to correlate the PTM information of the residues with these structures [80]. On the other hand, it is essential to concatenate the previous experimental evidence and construct a three-dimensional model for artificial intelligence learning.

## 3. Discussion

MDD is a pressure stress-mediated physiological disorder that often leads to reduced synaptic plasticity, in which the TrkB receptor, the starting point of the synaptic plasticity pathway, is an RTK distinct from 5-HT and glutamate receptors [11].Its large extracellular receptor and intracellular kinase structures, as well as the highly variable transmembrane helices and disordered sequences between the two, have led to the resolution of its activation state being mostly limited to the extracellular segment. Following AF2 catch all protein in one draft, AF-M evolved to be able to predict complex structures, allowing us to construct TrkB dimerization activation structures as well as complex structures with intracellular signal molecules. Adopting a divide and conquer algorithm, we first predicted the activation structure of the whole extracellular segment of TrkB binding to mBDNF and were able to rapidly perform homologous ligand-receptor pairing, providing three-dimensional information that can be used to design targeted agonists or antagonists that are highly specific to a given combination. For example, the design of mBDNF-mimetic microproteins targeting TrkB reduces the aberrant activation of TrkA and TrkC.

Transmembrane helices, due to their complex environment and their role as relay stations for signal cascades, often require careful design of membrane mimics for resolution. For unresolved transmembrane helix dimers, manual screening is required among numerous docking results. AF-M provides dimeric structures that are stable in the lipid raft environment and create pockets at the crossover that bind to novel antidepressants. This will strongly facilitate the design of drugs against RTKs, breaking the inherent impression that transmembrane helical drug targets exist only for GPCRs and ion channels with multiple transmembrane helices. It will also enhance the resolution of highly flexible dimeric transmembrane helices of RTKs under Cryo-EM and refine the transmembrane dimeric activation pattern of RTKs members in combination with MDS.

As a kinase segment that functions as a biochemical reaction, its potential chaperones exceed 300 species, and the complex conformation is currently difficult to capture due to the rapidly proceeding phosphorylation reaction [81]. Attempts were made to construct a complex pattern of TrkB kinase segment with three typical signal chaperones, and the binding site and pattern of PLCγ1 were highly similar to that of the resolved FGFR2 [31]. Unfortunately, however, the binding site of SHP2 presenting a self-repressed structure was designated as being in the disordered region due to the absence of PTM information. Rather, SHC1 successfully distinguished a binding mode with the PTB structural domain binding pTyr. Further iterations of AF-M are needed in the prediction of kinase-signal chaperones to predict the metastable binding process of proteins in biochemical reactions, such as researchers adding PTM information when uploading structures, marking key sites regulating metastable conformations, and uploading multiple dynamic conformations of the binding process to the database for artificial intelligence learning.

AF-M extends the prediction to multimers, giving us access to the underlying structures of molecular machine-protein complexes that dominate physiological functions. Such breakthroughs allow me to stand at the beginning of the next era of structural science, using deep learning-based artificial intelligence programs represented by AF2, combined with MDS, to build a dynamic signal pathway network containing structural information. In this network, the self-assembly of small molecules in all physiological and pathological processes can be demonstrated, and the operation of the entire pathway can be controlled by modifying and controlling a few key targets, which will enable drug design to shift from single-target to global [82]. In the foreseeable future, AI, a powerful assistant, will play an important role in breaking down difficult-to-treat diseases involving multiple mechanisms, such as MDD.

## 4. Materials and Methods

### 4.1 AF2 predicts the full-length structure of TrkB

To predict the full-length structure of TrkB, the first one was obtained directly from the AF Protein Structure Database (Last updated in Alphafold database, version 2022-06-01, created with the AF Monomer v2.0 pipeline). The second one is based on AF Monomer V2.2, manually inputting the full-length sequence of TrkB obtained from uniport. The third one is based on the monomer_casp14 program, which improves the average GDT of Monomer by about 0.1. AF2 was downloaded from github and run locally as described (https://github.com/deepmind/AF), using default parameters and the database version used: pdb _mmcif and uniport are 2022-08-03, the rest of the database is the default database [83].

### 4.2 AF-multimer predicts the structure of protein complexes

Human full-length sequences of TrkB, BDNF, SHP2, PLCγ1, and SHC were obtained from uniport, and sequences were selected for combination to construct multimers as needed, and the sequence combinations used are shown in the Supplementary Material. Run the AF-multimer program [37] and set ‘--model_preset=multimer’ to output the PDB file of the top 5 predicted complex structures sorted by PLDDT. All raw structures not shown are shown in the supplementary material.

### 4.3 Conformational optimization of drug small molecule drawing

The (2R,6R)-HNK, IHCH-7086 structures were drawn using ChemDraw, calling the Chem.AllChem module of RDKi (http://rdkit.org) using the Embed Molecule function using Experimental-Torsion Basic Knowledge Distance Geometry (ETKDG) algorithm [84] to generate 3D conformations based on the modified distance geometry algorithm, optimize and calculate the energy using MMFF94 force field [85], and finally select the lowest energy conformation as the initial conformation for docking.

### 4.4 Protein-drug molecule docking

The highest confidence 427-459 double alpha helix structure obtained by AF-M was taken, and Smina [86] was selected as the docking software for molecular docking with (2R,6R)-HNK and IHCH-7086. The positions of the active centers were as follows: X-center = -7.729, Y-center = 2.684, and Z-center = -6.868, where the approximation (exhaustiveness) of docking was 80, the box size of docking was 40 Å, and 80 Å conformations were generated each time, and the optimal conformations were selected for molecular dynamics simulations.

### 4.5 Analysis of protein-ligand interactions

Upload the pdb file of protein complex to PILP [87] for interaction analysis, set the A chain as the main chain, obtain the salt bond, hydrophobic bond, ππ bond, etc. generated with the interacting ligands, and visualize the output pse file with pymol.

### 4.6 Sequences alignment

Clustal Omega [88] is used to perform multiple sequence alignment, input the sequence information obtained from uniport, and set the output form as ClustalW with character counts. Visualizing the comparison results using Jalview [89]. The higher the similarity, the darker the color of the residues.

### 4.7 Molecular dynamics simulation

The conformation of ranked 1 transmembrane helical dimer coupled with (2R,6R)-HNK predicted by AF-M was added with 20% CHOL+80% DOPC as the membrane environment. Gromacs2019.6 [90] was chosen as the kinetic simulation software and amber14sb as the protein force field. Small molecules were used to produce topology files based on GAFF2 (Generation Amber Force Field) force field. The TIP3P water model was used to add TIP3P water model to the complex system to create a water box and add sodium ions to equilibrate the system. Under elastic simulation by Verlet and cg algorithms respectively, PME deals with electrostatic interactions and energy minimization using steepest descent method for maximum number of steps (50,000 steps). The Coulomb force cutoff distance and van der Waals radius cutoff distance are both 1.4 nm, and finally the system is equilibrated using the regular system (NVT) and isothermal isobaric system (NPT), and then the MDS simulations are performed for 100 ns at room temperature and pressure. In the MDS simulations, the hydrogen bonds are constrained by the LINCS algorithm with an integration step of 2 fs. The Particle-mesh Ewald (PME) method is calculated with a cutoff value of 1.2 nm, and the non-bond interaction cutoff value is set to 10 Å. The simulation temperature is controlled by the V-rescale temperature coupling method at 300 K, and the pressure is controlled by the Berendsen method at 1 Å. NVT and NPT equilibrium simulations were performed at 300 K for 30 ps, and finally, the finished MDS simulations were performed for 50 ns. The root mean square deviation (RMSD) was used to observe the local site variation during the simulation (the cut-off point was set to 0.2). The radius of gyration (Rg, radius of gyration) is used to evaluate the tightness of the structure of the system. The root mean square function (RMSF) is used to observe the local site metastability of the system during the simulation, and the solvent accessible surface area (SASA) is used to observe the size of the solvent accessible surface area of the complex during the simulation.

### 4.8 Binding free energy calculation for proteins and small molecules

The MDS trajectory is used to calculate the binding free energy by the following equation:

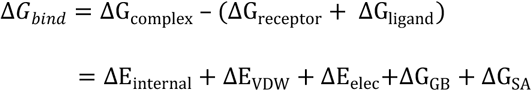

In the above equations, ΔE_internal_ internal represents internal energy, ΔE_VDW_ represents van der Waals interaction and ΔE_elec_ represents electrostatic interaction. The internal energy includes the bond energy (E_bond_), angular energy (E_angle_), and torsion energy (E_torsion_); ΔG_GB_ and ΔG_GA_ are collectively referred to as the solvation free energy. Among them, G_GB_ is the polar solvation free energy and G_SA_ is the non-polar solvation free energy. For ΔG_GB_, the GB model is used for calculation (igb = 8). The nonpolar solvation free energy (Δ*G*_*SA*_) is calculated based on the product of surface tension (γ) and solvent accessible surface area (SA), GSA= 0.0072 × SASA [91]. The entropy change is neglected in this study due to the high computational resources and low precision. this study is neglected. This algorithm is implemented by Gmx_MMPBSA [92].

## Author Contributions

Conceptualization, C.Q., X.X and H.Y.; methodology, C.Q. and H.Y.; software, P.L. and Y.L.; validation, P.L. and X.X.; formal analysis, B.C.,D.W. and W.A.; investigation, C.Q. and H.Y.; resources, M.C.,L.W.,T.A. and J.X.; data curation, X.X.; writing–original draft preparation, C.Q. and H.Y.; writing–review and editing, X.L. and F.X.; visualization, C.Q.,P.L. and X.X.; supervision, B.L., X.L. and F.X.; project administration, B.L.,X.L. and F.X.; funding acquisition, X.L. and F.X. All authors have read and agreed to the published version of the manuscript.

