## Supplementary Materials for "Reconstruction of TrkB complex assemblies and localizing antidepressant targets using Artificial Intelligence"

**Table of content:**

**Figure S1-S3: Alphafold 2 predicted TrkB monomers and dimers**

**Figure S4-S5: Details of the contact between the extracellular segment of TrkB and mBDNF**

**Figure S6-S7: MDs data of TrkB transmembrane helix, TrkB docking with ADs**

**Figure S8-S12: Multiple possible posture of TrkB assemblies**

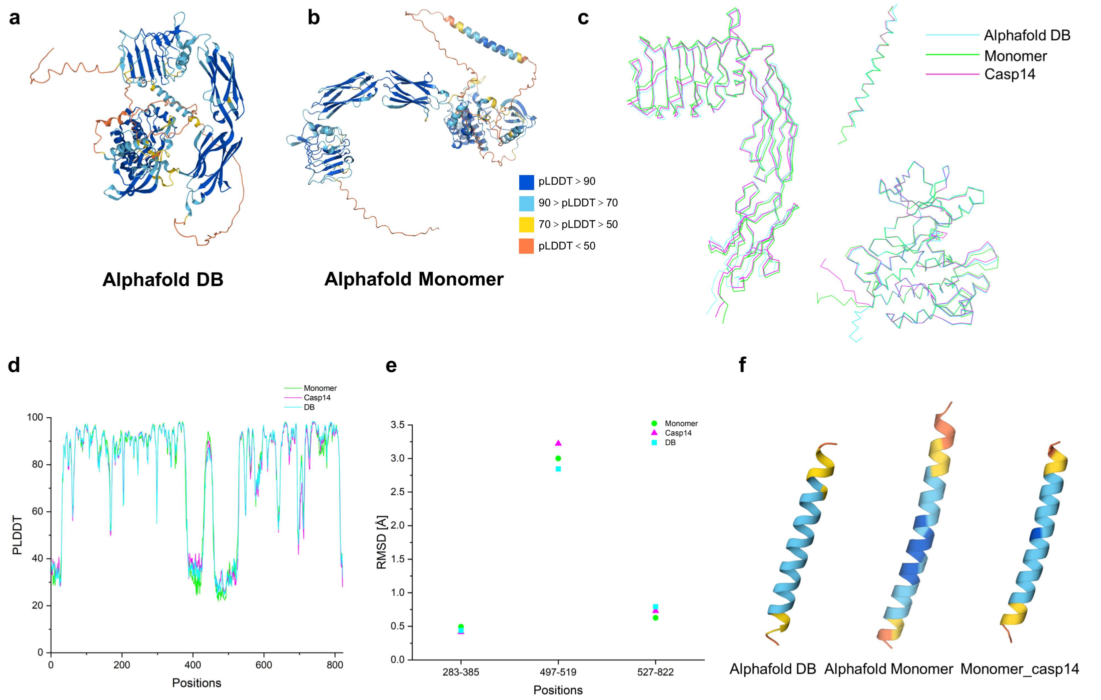


**Fig S1**. (**A**) Full-length structure of TrkB monomer obtained from Alphafold Database. PLDDT is distinguished by different colors (same later). (**B**) Full-length structure of TrkB monomer predicted by Alphafold Monomer V2.2. (**C**) Overlay of structures of DB, Monomer, and Casp14 fixed structures shown as ribbons. (**D**) PLDDT distribution of full-length residues of DB, Monomer, Casp14. (E) RMSD values of DB, Monomer, Casp14.

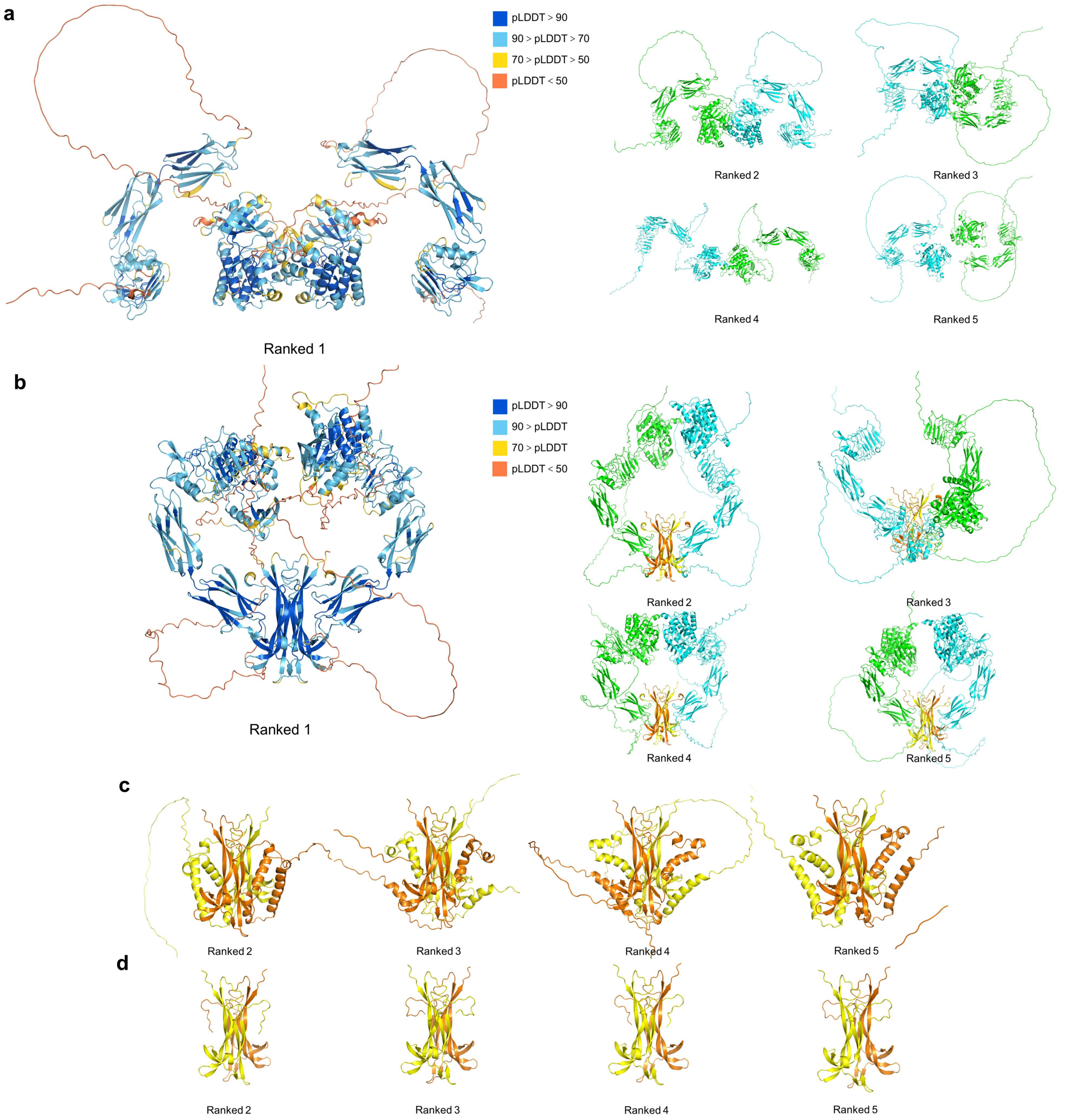


**Fig S2.** (**A**) Top five possible outcomes of AF-M prediction of TrkB full-length dimer structure. (**B**) Top five possible outcomes of AF-M predicted TrkB full-length dimer bound mBDNF structure. (**C**) AF-M predicts the ranked 2-4 of proBDNF dimer structure. (**D**) AF-M predicts the ranked 2-4 of mBDNF dimer structure.

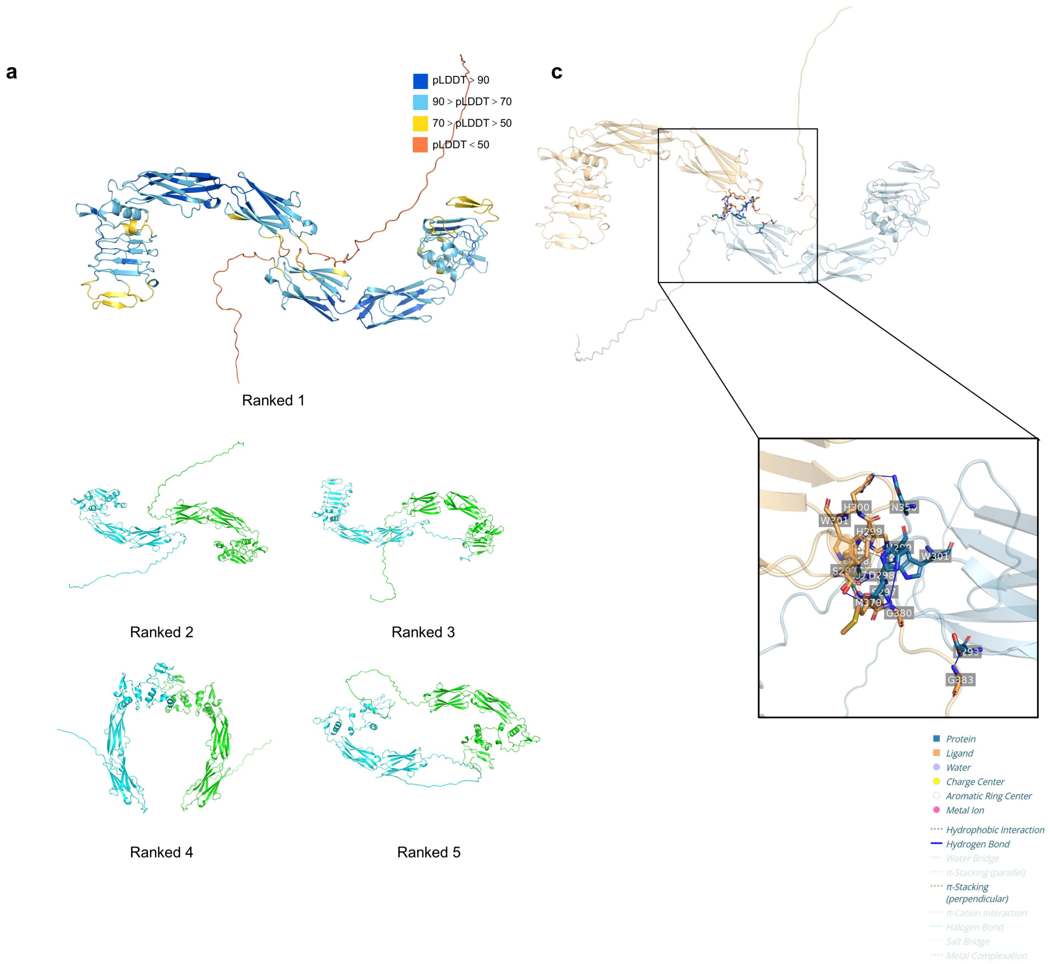


**Fig S3.** (**A**) AF-M prediction of the top five possible outcomes of the TrkB extracellular segment dimer structure. (**B**) Residue interactions between the two monomers of Ranked 1 are shown.

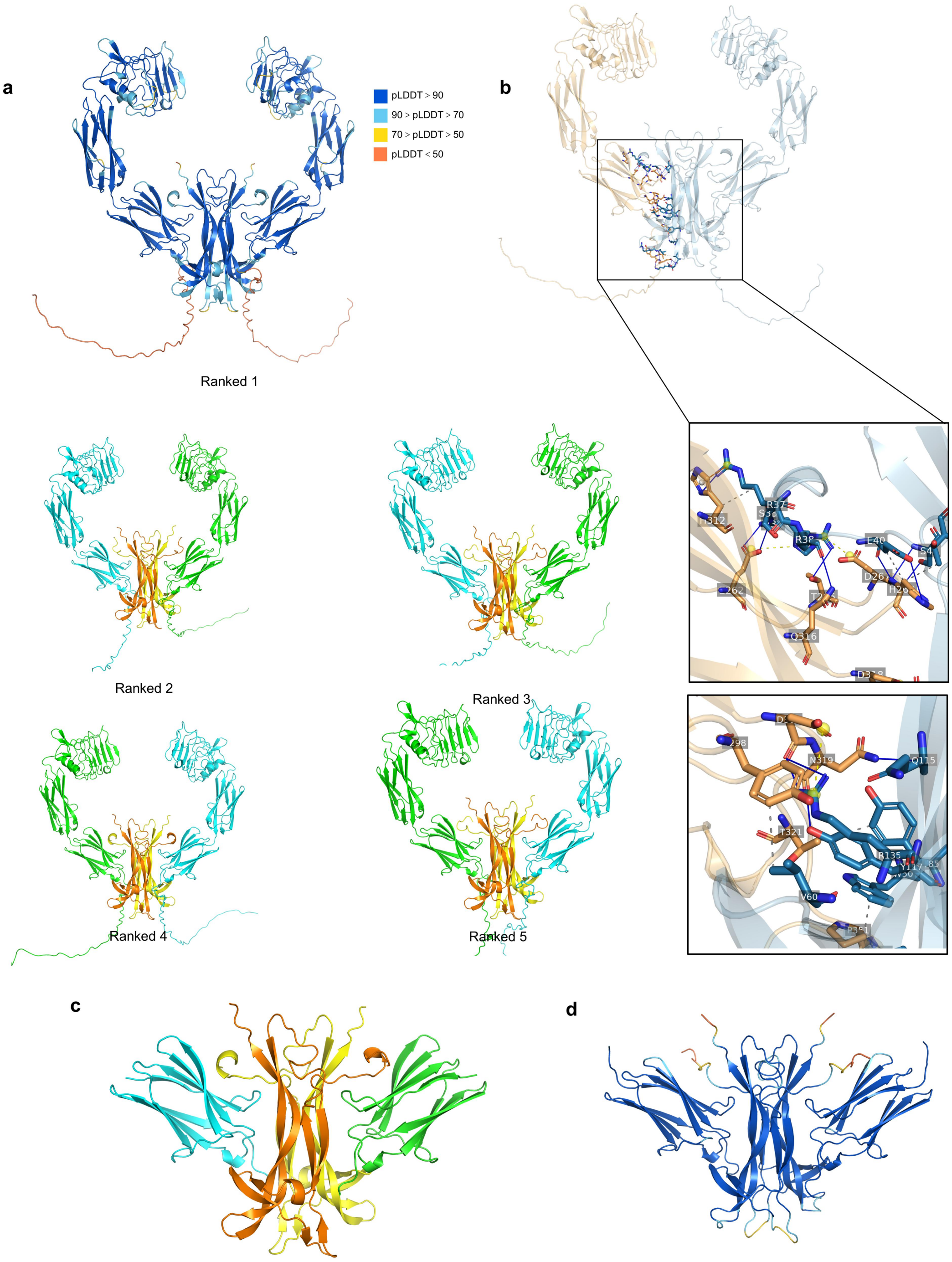


**Fig S4.** (**A**) AF-M prediction of the top five possible outcomes of TrkB extracellular segment dimer binding to mBDNF structure. (**B**) Residue interactions between the Ranked 1 monomeric extracellular segment and mBDNF are shown. (**C**) Structure of the TrkB domain3-bound mBDNF complex predicted by AF-M. (**D**) Structure of the AF-M predicted complex of TrkC domain3 binding NTF-3.

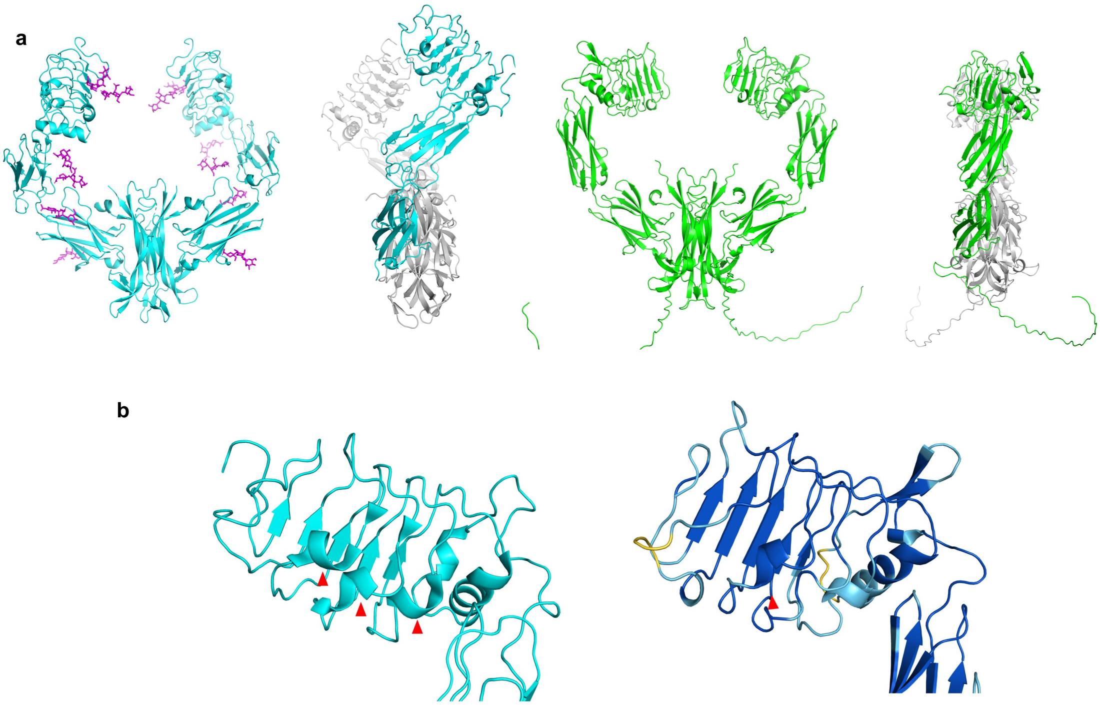


**Fig S5.** (**A**) Structure of the experimental complex of TrkA extracellular segment with NGF (PDB: 2IFG), Asn-linked glycosylation is labeled in purple. Side view species Domain1~3 show a Z-fold (left). AF-M predicts the TrkB extracellular segment dimer binding mBDNF activation structure, the angle between Domain1~3 in side view is smaller than TrkA and lies in the same plane (right). (**B**) The experimental complex of TrkA extracellular segment with NGF has a total of three incomplete helices at the articulation of Domain1, LRR (left). AF-M predicts the TrkB extracellular segment dimer binding mBDNF structure of Doamin1, LRR with only one incomplete helix at the articulation and the overall structure of Domain 1 is flatter and rounder (right).

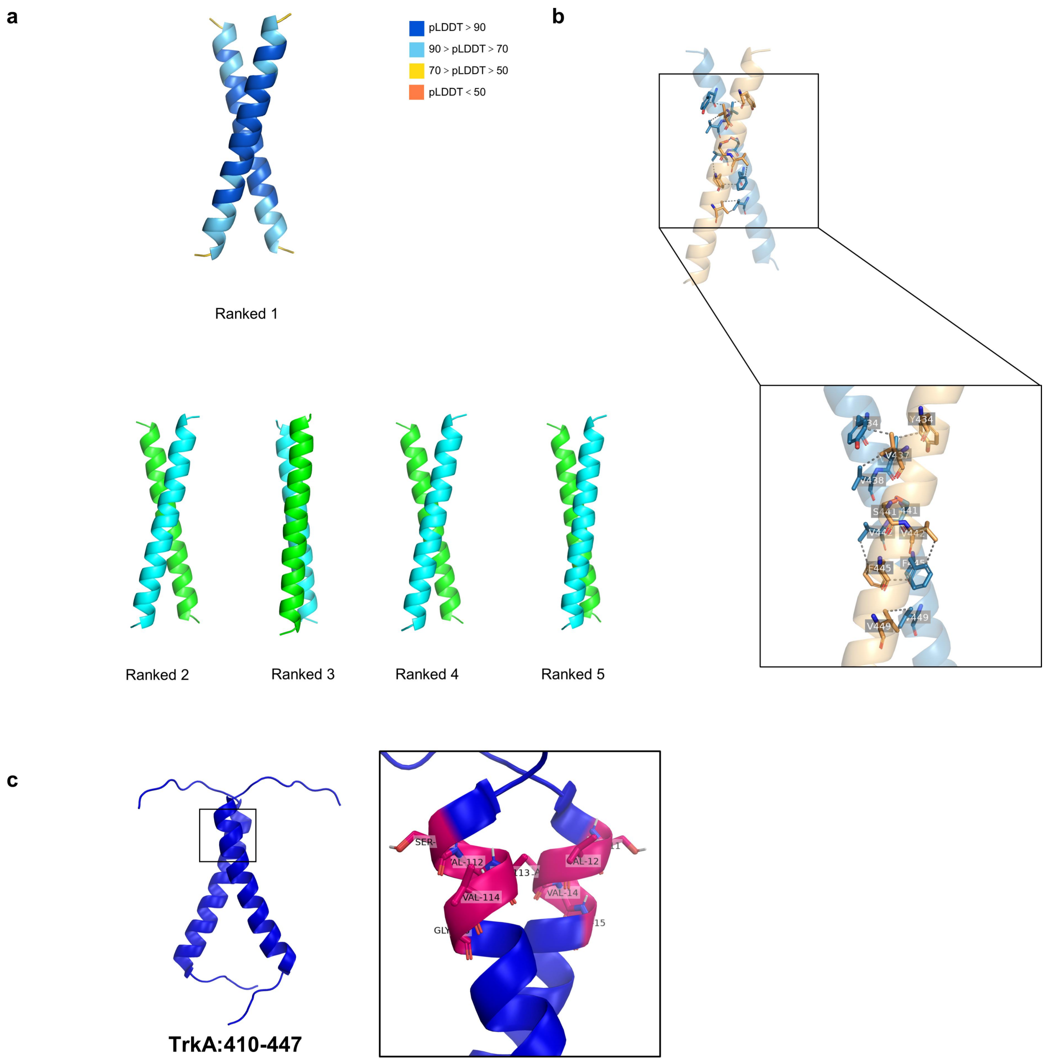


**Fig S6.** (**A**) AF-M prediction of the top five possible outcomes for the dimeric structure of the transmembrane segment of TrkB. (**B**) Residue interactions between two monomers of Ranked 1 are shown. (**C**) Experimental structure of TrkA transmembrane helical dimer with cross-residue SVAVG shown in red stick form.

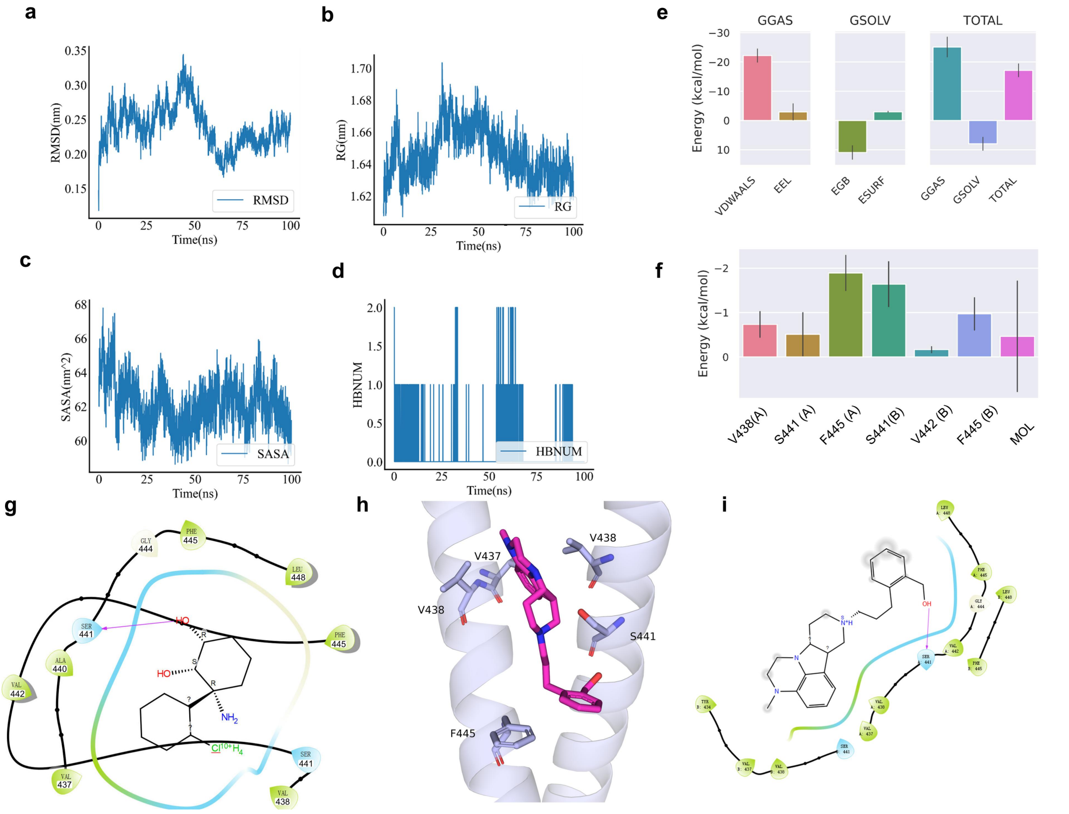


Fig S7. Molecular dynamics simulation after transmembrane helix docking of (2R,6R)-HNK. (**A**) RMSD values of the complexes during 100 ns simulation, fluctuating from 50-60 ns and entering the steady state after 75 ns. (**B**) The radius of gyration (RG) values of the complex during the 100ns simulation, with a small peak at 0ns, starting to reach a second peak when entering 50ns, and finally entering the stable interval after 75-100ns. (**C**) The solvent accessible surface area (SASA) of the complex during the 100ns simulation, with a decreasing trend of SASA fluctuation. (**D**) Hydrogen bonding change curve (HBNUM) of the complex during the 100ns simulation, the complex has 0-2 connections of hydrogen bonds in the steady state. (**E**) The decomposition of the free energy properties between (2R,6R)-HNK and the transmembrane helix. the TOTAL free energy is -17.14 kcal/mol, indicating that the two have binding possibilities. (**F**) 2R,6R)-HNK degradation map with free energy residues between transmembrane helices, V438(A), S441(A), F445(A), S441(B), V442(B), F455(B) were favorable for binding. (**G**) Pose fingerprint of transmembrane helix dimer with (2R,6R)-HNK. Blue is polar amino acid, green is hydrophobic amino acid. The purple arrows represent the direction of the interaction force, pointing from the donor to the acceptor. (**H**) Complex conformation before and after MD simulation, with the transmembrane helix tightened inward and (2R,6R)-HNK moving to where the helices intersect. (**I**) Pose fingerprint of the transmembrane helix dimer IHCH-7086.

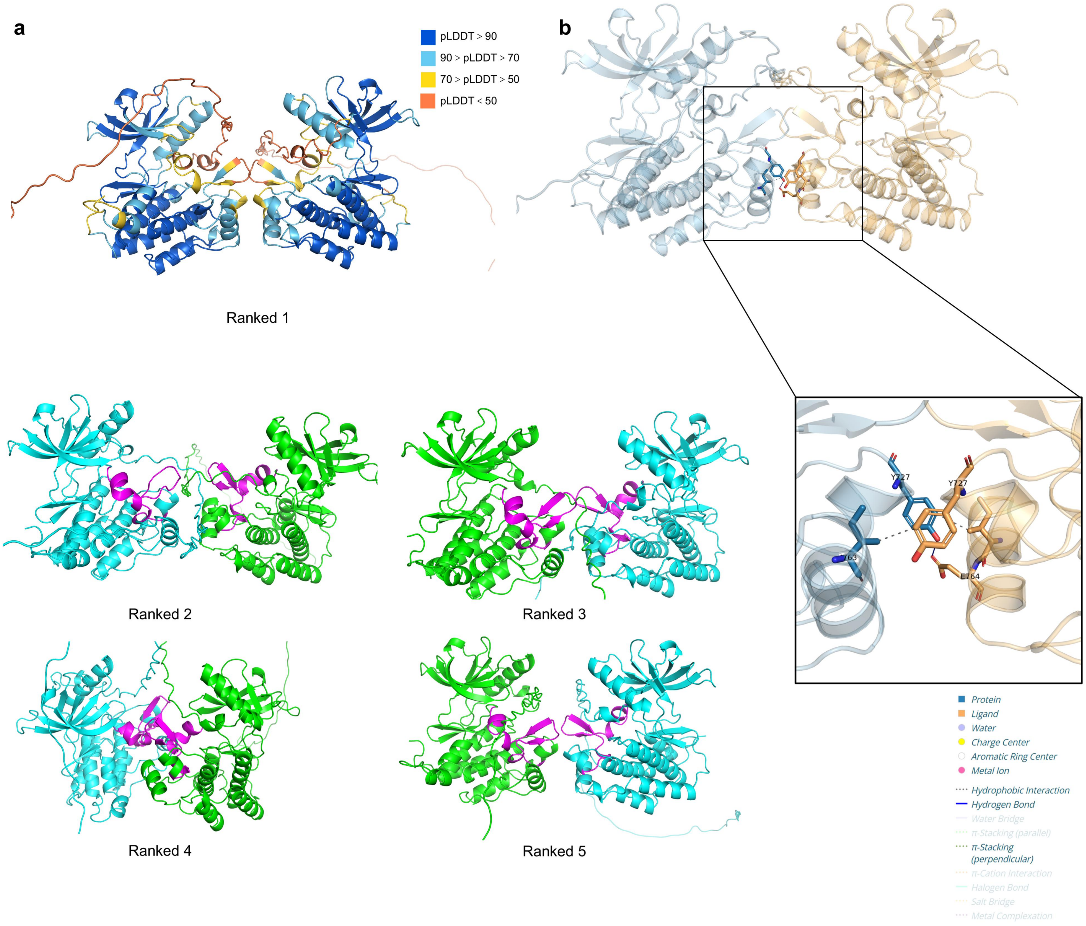


**Fig S8.** (**A**) AF-M prediction of the top five possible outcomes for the dimeric structure of the intracellular segment of TrkB. (**B**) Residue interactions between two monomers of Ranked 1 are shown.

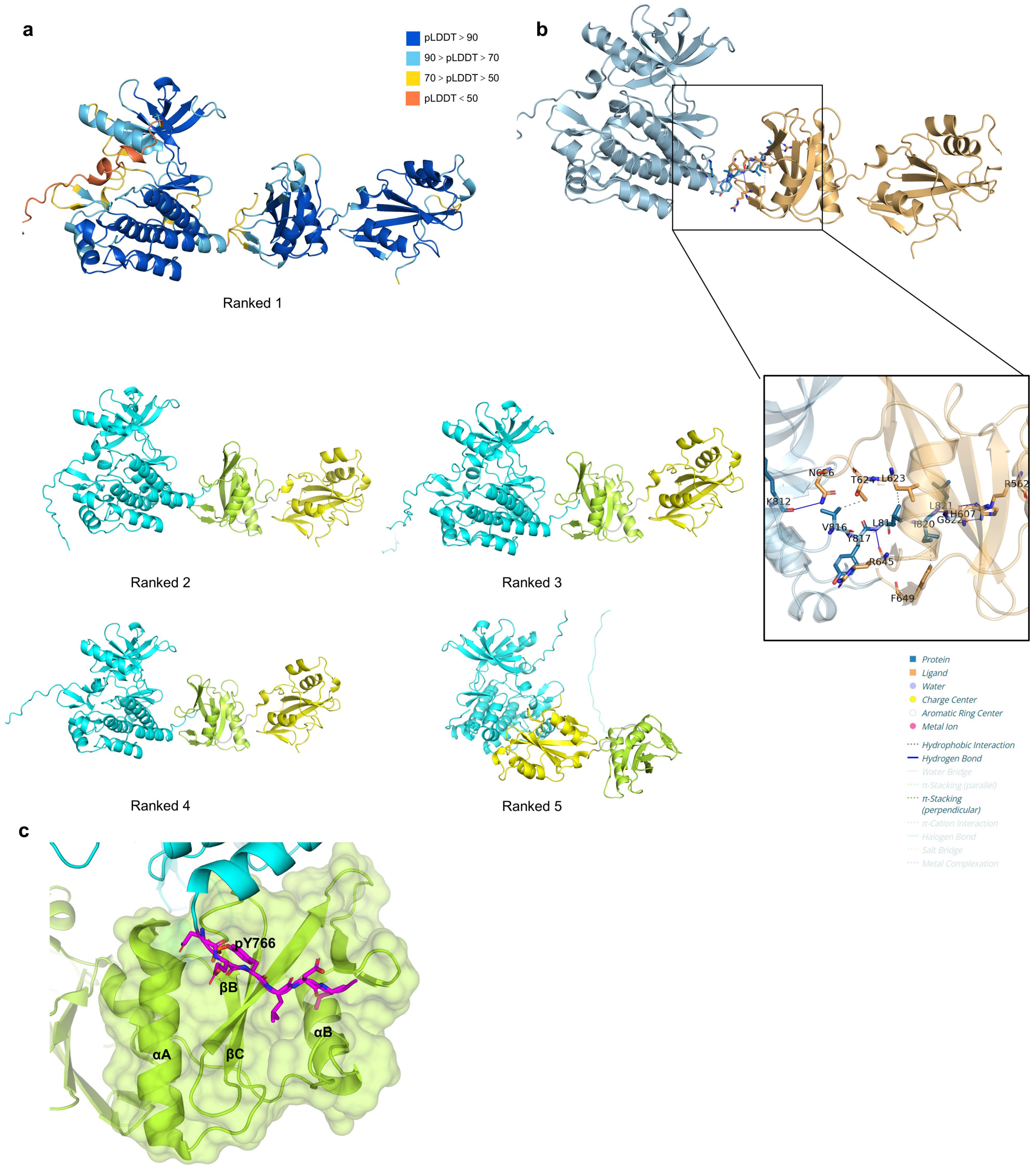


**Fig S9.** (**A**) AF-M prediction of the top five possible outcomes of the intracellular segment of TrkB binding to the PLCγ1 NSH2-CSH2 structure. (**B**) Residue interactions between two monomers of Ranked 1 are shown. (**C**) Detail diagram of contact residues of the experimental structure of FGFR1 in complex with PLCγ1 NSH2-CSH2.

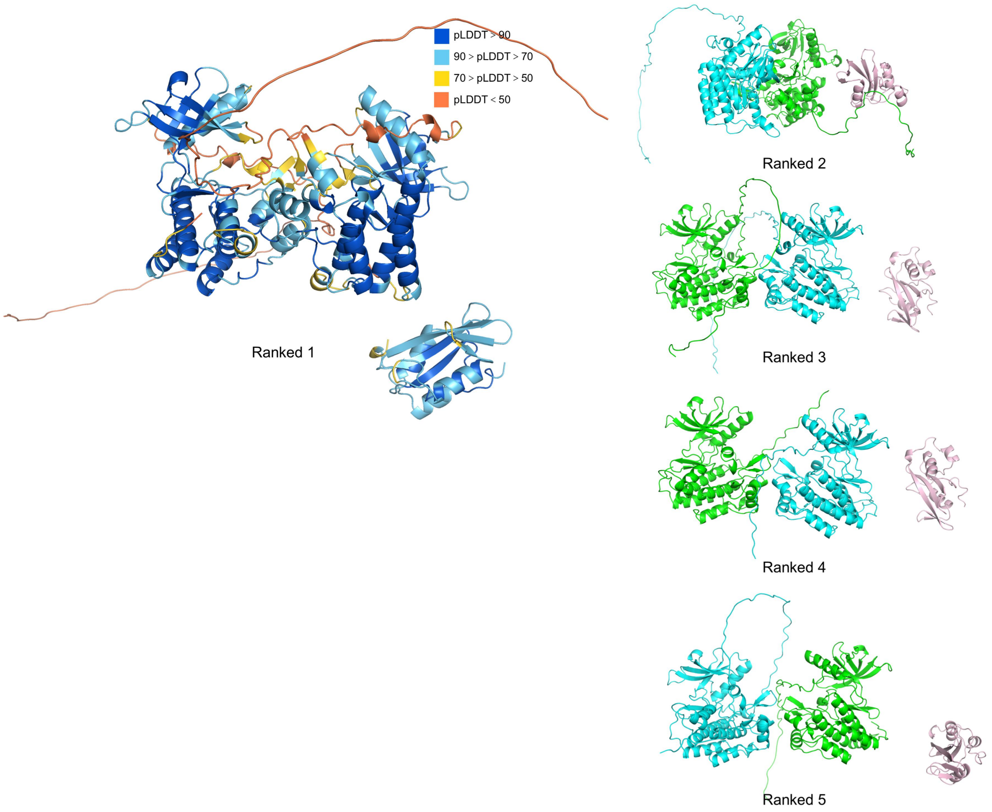


**Fig S10.** AF-M predicts the top five possible outcomes of TrkB intracellular segment binding to PLCγ1 CSH2.

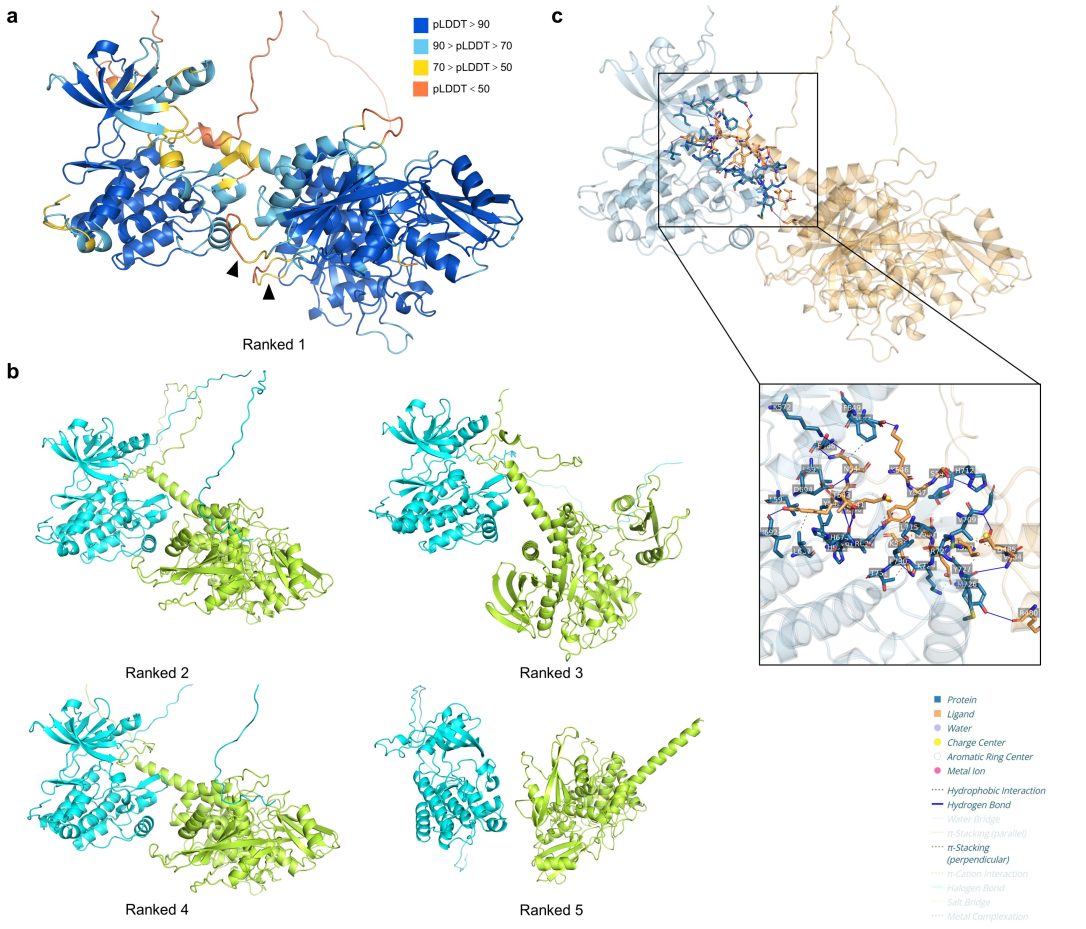


**Fig S11.** (**A**) AF-M prediction of the top five possible outcomes of TrkB intracellular segment binding to SHP2. (**B**) Residue interactions between two monomers of Ranked 1 are shown.

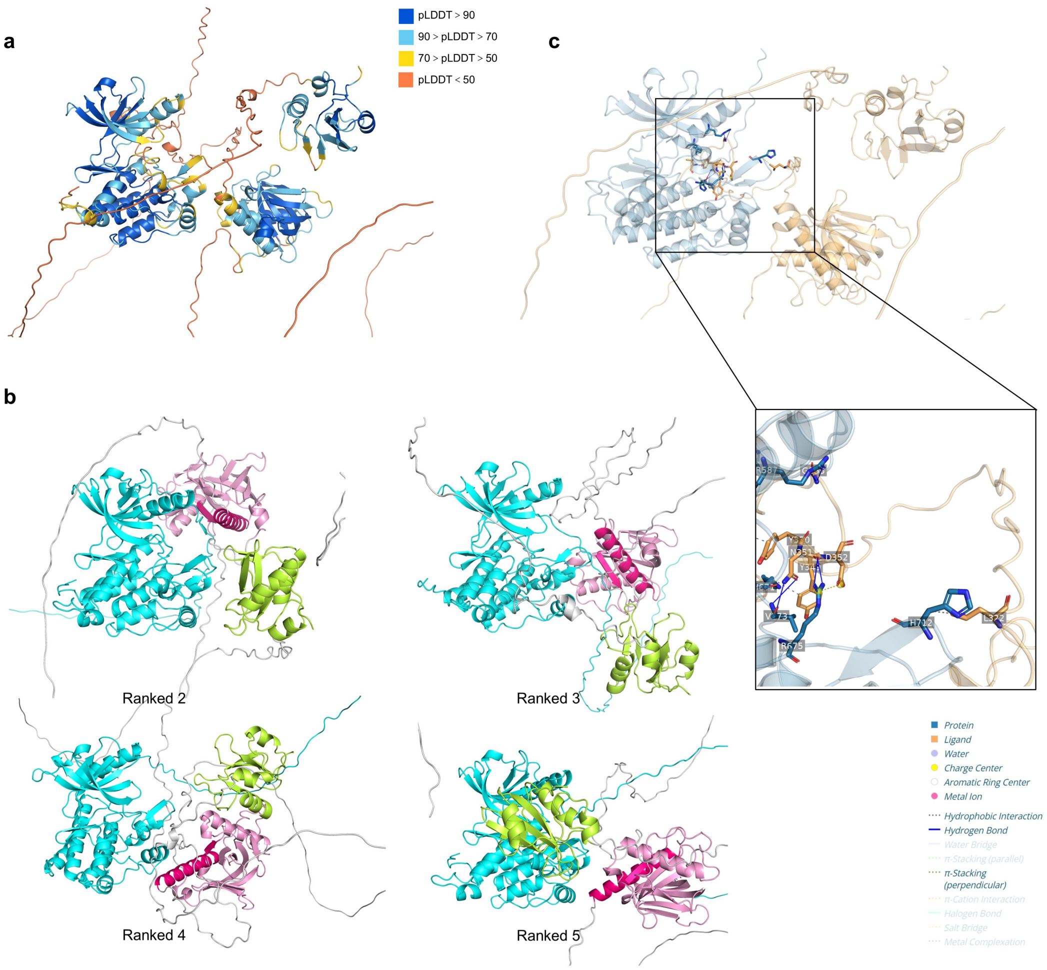


**Fig S12.** (**A**) AF-M predicts the top five possible outcomes of TrkB intracellular segment binding to SHC1. (**B**) Residue interactions between two monomers of Ranked 1 are shown.
